# Predicting Individualized Functional Topography in Developmental Prosopagnosia

**DOI:** 10.64898/2026.03.18.712539

**Authors:** Ian Abenes, Guo Jiahui

## Abstract

Functional localizer scans have long served as the classic method for mapping individualized functional topographies, but they require dedicated scan time and can be difficult to implement in neuropsychological populations. Previous work has shown that individualized functional topographies can be estimated with high fidelity in typical participants using hyperalignment, but it remains unknown whether this approach generalizes to populations with functional deficits. Here, we tested this question in developmental prosopagnosia (DP), a neuropsychological condition characterized by severe face recognition impairments. Using two independent datasets that included both DP and control participants, we estimated individualized category-selective functional topographies from independent participants using hyperalignment derived from either a task-based scan or a naturalistic movie-viewing scan. Across datasets, whole-brain correlations and searchlight analyses showed that predicted topographies were highly similar to topographies estimated from participants’ own localizer data, especially in cortical areas with strong category-selective responses. Hyperalignment successfully recovered idiosyncratic features of category-selective topographies and consistently outperformed anatomical alignment. Importantly, predictions generalized across groups, such that individualized topographies in DPs could be estimated from control participants and vice versa. In addition, predicted topographies preserved the reduced face selectivity in DPs previously reported in the literature. These findings support a hyperalignment-based framework for estimating individualized functional topographies in neuropsychological populations without requiring separate localizer scans, and provide a foundation for integrating existing datasets to study the underlying neural basis in DP and other atypical populations.

## Introduction

Developmental prosopagnosia (DP) is characterized by great difficulty in face recognition despite normal low-level vision, intelligence, and no history of brain damage (Susilo & Duchaine, 2013; Behrmann et al., 2016). Significant behavioral and neural heterogeneity has been found in DPs, with reports of atypical neural tuning and response patterns within and beyond the face-processing network (DeGutis et al., 2024; Manippa et al., 2023). Furthermore, neuroimaging studies have demonstrated substantial individual differences in the activation, spatial extent, and anatomical location of category-selective topographies in typical participants (Zhen et al., 2015). This substantial topographical variability, combined with neural abnormalities in DPs, makes this group particularly valuable for evaluating methods that estimate individualized functional topography in populations with atypical category-selective responses.

Functional localizer scans have long served as a standard and widely adopted method for mapping individualized functional topographies. By contrasting well-controlled stimulus categories, localizers provide interpretable and reliable estimates of category-selective regions at the individual level (e.g., Kanwisher et al., 1997; Saxe et al., 2006, Fedorenko et al., 2011). However, because each localizer is typically optimized to probe a limited set of functional contrasts, multiple dedicated scans are often required to characterize different functional systems. In practice, researchers may need to balance the number of localizer runs and stimulus categories against constraints of scan time and resources. In addition, extended or repeated localizer scans can place sustained demands on participants’ attention, which may limit their feasibility in certain populations, including children and neuropsychological or clinical groups. These considerations can pose challenges for studying special populations at scale. At the same time, individual laboratories are often limited in their ability to recruit large samples from special populations, and many existing studies consequently rely on modest sample sizes. Integrating datasets collected across sites, scanners, and research goals, therefore, presents an important opportunity to increase statistical power and maximize the value of existing data. Nevertheless, variability in localizer design and implementation across studies introduces heterogeneity that can complicate direct dataset integration using conventional functional mapping approaches.

These practical considerations motivate the need for an approach that can estimate individualized functional topography with high fidelity while maintaining flexibility in experimental design and data collection, and effectively leveraging existing datasets with substantial variability. Previous work has demonstrated that individualized functional topographies can be estimated with high fidelity using hyperalignment (Guntupalli et al., 2016; Jiahui et al., 2020). Using either response-based (Jiahui et al., 2020) or connectivity-based hyperalignment (Jiahui et al., 2023), functional topography in a new participant can be predicted using only movie data from that participant together with movie and localizer data from a group of independent participants. In this framework, movie data from the new participant are used to derive transformation matrices that align the target participant and each individual in the independent group, and these transformation matrices are then applied to localizer data of each individual from the independent group to project localizer runs into the target participant’s space. The estimated topography for the target participant was obtained by averaging the predicted topographies across all individuals in the independent group. This approach has been validated across multiple datasets and different category-selective maps.

The primary goal of the present study is to establish a framework for estimating individualized functional topographies in individuals with DP using hyperalignment with independent participants based on either a separate task or a naturalistic movie. To address this question, we first used a previously collected dataset comprising one of the largest samples in the DP literature (Jiahui et al., 2018). This dataset included individuals with DP and matched control participants, each scanned with a category-selective dynamic functional localizer (Pitcher et al., 2011) as well as an independent task. Either the independent localizer runs or the task scans were used to derive transformation matrices, which were then applied to held-out localizer runs from independent participants. We examined estimation both within groups (control-to-control, DP-to-DP) and across groups (control-to-DP, DP-to-control) to evaluate performance across different populations. To further test generalizability with a naturalistic design, we additionally analyzed the only publicly available dataset with DP that includes naturalistic movie-viewing scans (Noad et al., 2024). In this dataset, participants watched clips from Game of Thrones and were also scanned with a brief static category-selective localizer. Finally, to examine whether predicted topographies preserve the differences in category selectivity between groups in previous analyses, we conducted region-of-interest (ROI) analyses (Norman-Haignere et al., 2016; Jiahui et al., 2018) using the estimated functional maps. We found that individualized functional topographies were successfully estimated with high fidelity using independent scans and participants from either the same or different neuropsychological and control groups, and that these estimates preserved functional deficits previously reported in DPs (Manippa et al., 2023).

This study establishes a framework for estimating individualized functional topography from independent task-based and naturalistic scans in a neuropsychological population. This framework is ready to be generalized across experimental contexts and data sources, and provides a foundation for future large-scale studies that integrate heterogeneous datasets to investigate the neural basis of neuropsychological conditions across different types of functional areas and networks.

## Results

To investigate whether individualized functional topographies can be estimated with high fidelity in neuropsychological populations with hyperalignment from independent individuals and scans, we used two independent datasets with DPs (the Dartmouth dataset and the GoT dataset). Both datasets also include a matched group of control participants and provide functional localizer scans, which makes it possible to compare the topography estimated from others using independent data with the topography estimated from the target participant’s own localizer scans. Beyond localizer scans, the Dartmouth dataset includes an independent attentional modulation task, and the GoT dataset includes a naturalistic movie scan in which participants watched clips from the TV series Game of Thrones.

### Hyperalignment enables high-fidelity estimations of individualized topography in DPs

To demonstrate whether individualized predictions can be achieved with high fidelity in DP, we first focused on the localizer data in the Dartmouth dataset. We split the five runs into training and testing data following a leave-one-run-out procedure. For each pair of participants, transformation matrices were derived based on the training data using hyperalignment and applied to the left-out localizer run of the source participant to transform the data into the target participant’s space (Jiahui et al., 2020 & 2023). The predicted localizer runs were fit with a GLM, and the contrast maps were averaged across all source participants to derive the predicted topography for the target participant. We correlated topographies estimated from the target participant’s own localizer runs and from other source participants’ hyperaligned localizer runs to measure the prediction performance.

Both response hyperalignment (RHA), which was based on response time series, and connectivity hyperalignment (CHA), which was based on functional connectivity profiles, were used in this analysis. We used both within- and between-group predictions across DP and control participants. In the within-group predictions, both source and target participants were from the same group (i.e., the control participants’ group), and in the between-group predictions, the source and target participants were from two different groups (i.e., one from the control and one from the DP group). We found that in all four scenarios, hyperalignment outperformed traditional anatomical alignment (AA) (all *p* < 0.001, Bonferroni-corrected; Figures 1 & 2; also see Supplementary Materials Table 1) in predicting individualized category-selective topographies. The hyperalignment predicted performance was comparable to or exceeded the noise ceiling calculated using Cronbach’s alpha, based on contrast maps derived from individual localizer runs for each source participant (Figure 2). Cronbach’s alpha measured the mean reliability of the source localizer data.

**Figure 1.**
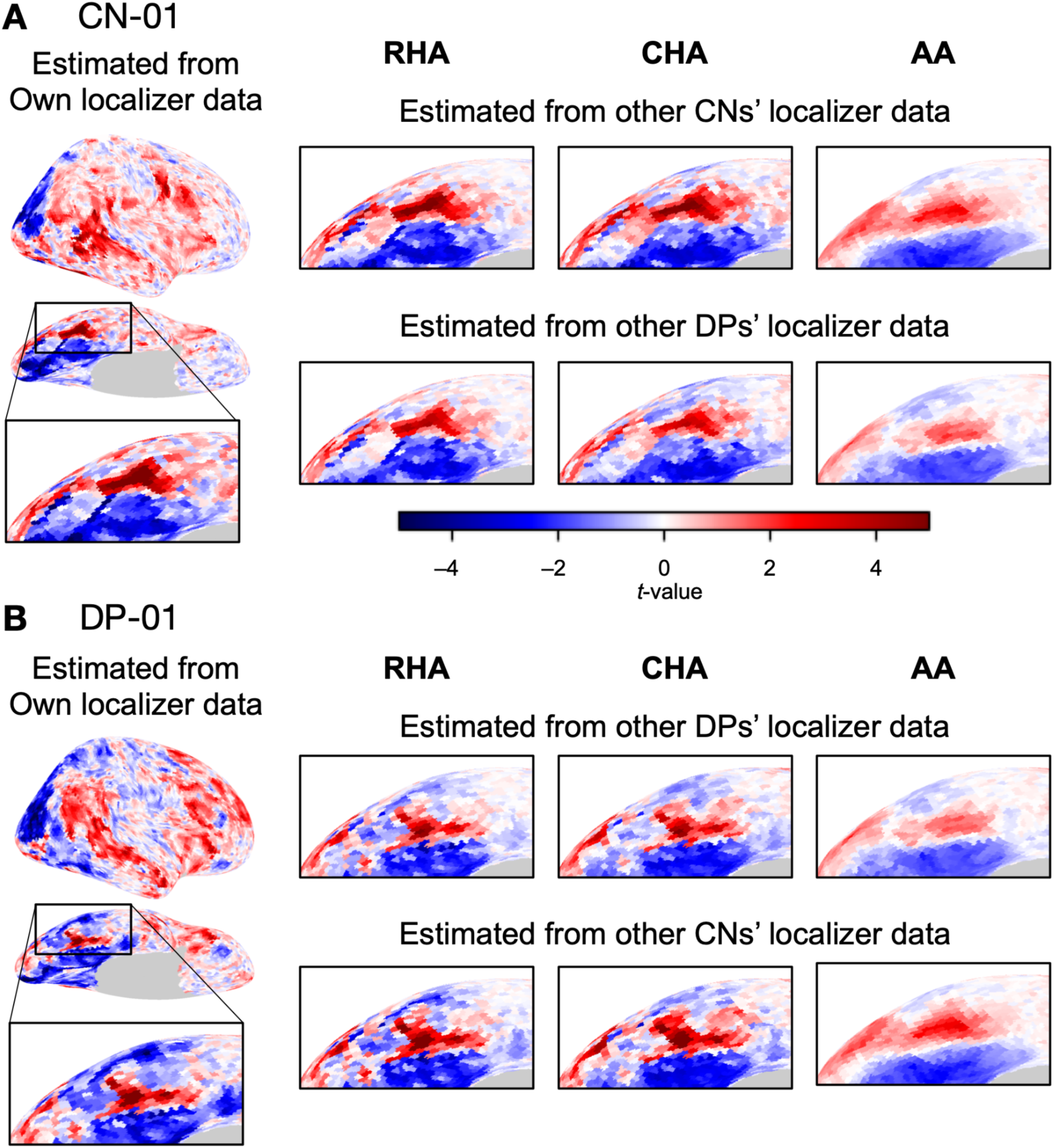
Sample contrast maps and enlarged views of the ventral temporal cortex of control and DP participants. Face-selective contrast maps (faces-vs-all) were estimated using localizer runs from the Dartmouth dataset. **(A)** The estimated face-selective contrast maps for a sample control participant, with expanded views of the right ventral temporal cortex. The left-most plot shows the map estimated from that control participant’s own localizer data. The grid shows contrast maps estimated from other participants using either response hyperalignment (RHA), connectivity hyperalignment (CHA), or anatomical alignment (AA). The top row shows maps estimated from other control participants, while the bottom row shows maps estimated from other DP participants. **(B)** The estimated face-selective contrast maps for a DP participant. As in **(A)**, the left-most plot shows the map estimated from the DP’s own localizer data, while the grid shows maps estimated from other participants. The top row shows estimations from other DP participants, while the bottom row shows estimations from control participants.

**Figure 2.**
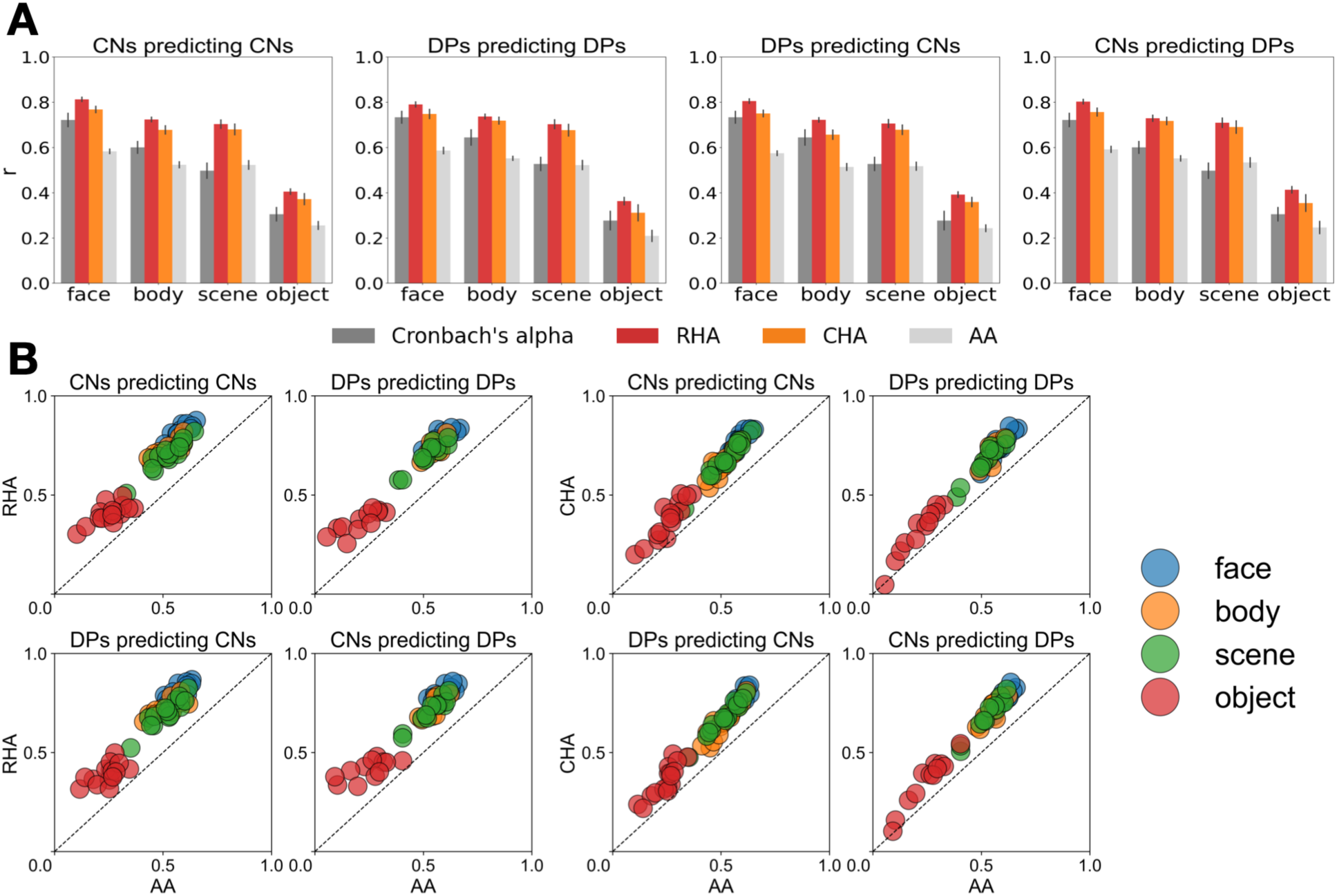
Correlations between contrast maps estimated from participants’ own localizer runs and from other participants’ localizer data. **(A)** Cronbach’s alpha values and mean whole-brain correlations across the four categories in the Dartmouth dataset. Each plot shows the Cronbach’s alpha values for each category (dark gray) and mean correlations between a participant’s contrast map and their predicted map from hyperalignment (RHA and CHA, red and orange, respectively) or anatomical alignment (AA, light gray). Cronbach’s alpha was calculated based on data from the source group for each prediction. From left to right, the first two plots show within-group comparisons (Controls to Controls, DPs to DPs), while the last two plots show across-group comparisons (DPs to Controls, Controls to DPs). In the across-group comparisons, control participants’ data were hyperaligned to DP participants’ space (and vice versa). In all plots, hyperalignment predicted results (RHA and CHA) are significantly better than AA predicted results (*p* < 0.001, Bonferroni corrected). Error bars stand for ±1 SE. **(B)** Scatter plots of RHA or CHA predicted performance versus AA predicted performance for each individual. Each dot represents a participant’s correlation coefficient between maps estimated using their own localizer data and using RHA or CHA (y-axis) in one of the four categories, plotted relative to the correlation coefficient using AA (x-axis). The two left-most scatter plots correspond to RHA, while the two right-most plots correspond to CHA.

Consistent with our previous results (Jiahui et al., 2020 & 2023), the predicted topographies successfully recovered the individualized size, locations, and even shape features of the target participant’s topographies estimated from their own localizer runs in the control participants group (Figure 1A). By contrast, predicted topographies based on anatomical alignment were blurry and smoothed, losing the idiosyncratic details of the individualized functional topographies and being essentially quite similar across individual target participants, as they were the direct average of topographies of the source participants. Prediction performance from RHA slightly outperformed CHA in a few categories (Supplementary Materials Table 1), but both RHA and CHA were significantly superior to AA, and both preserved idiosyncratic features in the predicted topographies, generating predicted individualized topographies with high fidelity.

Similarly, DPs showed a comparable within-group prediction performance, demonstrating successful prediction of a new DP’s category-selective topography based on other DPs (Figure 2). Most importantly and surprisingly, we found high-fidelity predictions in the across-group analysis, suggesting that a new DP’s individualized category-selective topographies can be estimated with superior performance from a group of typical participants’ localizer runs using hyperalignment and vice versa (Figures 1 & 2).

To investigate the brain areas that showed superior performance in prediction, we used a searchlight analysis with a 15mm radius to compare the topographies predicted from other participants using hyperalignment or AA and from their own localizer runs. We also calculated Cronbach’s alpha for each searchlight across localizer runs of the source participant’s group. Consistent with prior findings (Jiahui et al., 2020 & 2023), areas in the lateral occipital cortex (LOC), ventral temporal cortex (VT), superior temporal sulcus (STS), and frontal regions showed higher reliability (*r* > 0.8) in both groups. In all prediction scenarios, both RHA and CHA predicted topographies showed strong correlations with participants’ localizer-based topographies (*r* > 0.8) in broad areas, such as LOC and STS, especially in areas involved in face-selective processing (Figure 3). These areas demonstrated estimations that were close to or exceeded the reliability values, indicating high-fidelity prediction performance. On the other hand, prediction performance with AA was more right-hemisphere lateralized, with much weaker performance in nearly every part of the cortex. The right hemisphere lateralization in AA suggested high idiosyncratic localizations in the left hemisphere for face processing, and the weak overall performance across the cortex in AA was consistent with the whole-brain analysis in Figure 2. For the other categories, we found that areas involved in processing information of the preferred category (e.g., body-selective areas) showed stronger reliability and better prediction performance, similar to the face category. Additionally, for all categories, AA consistently showed weaker prediction performance across the entire cortex (Supplementary Figure S3).

**Figure 3.**
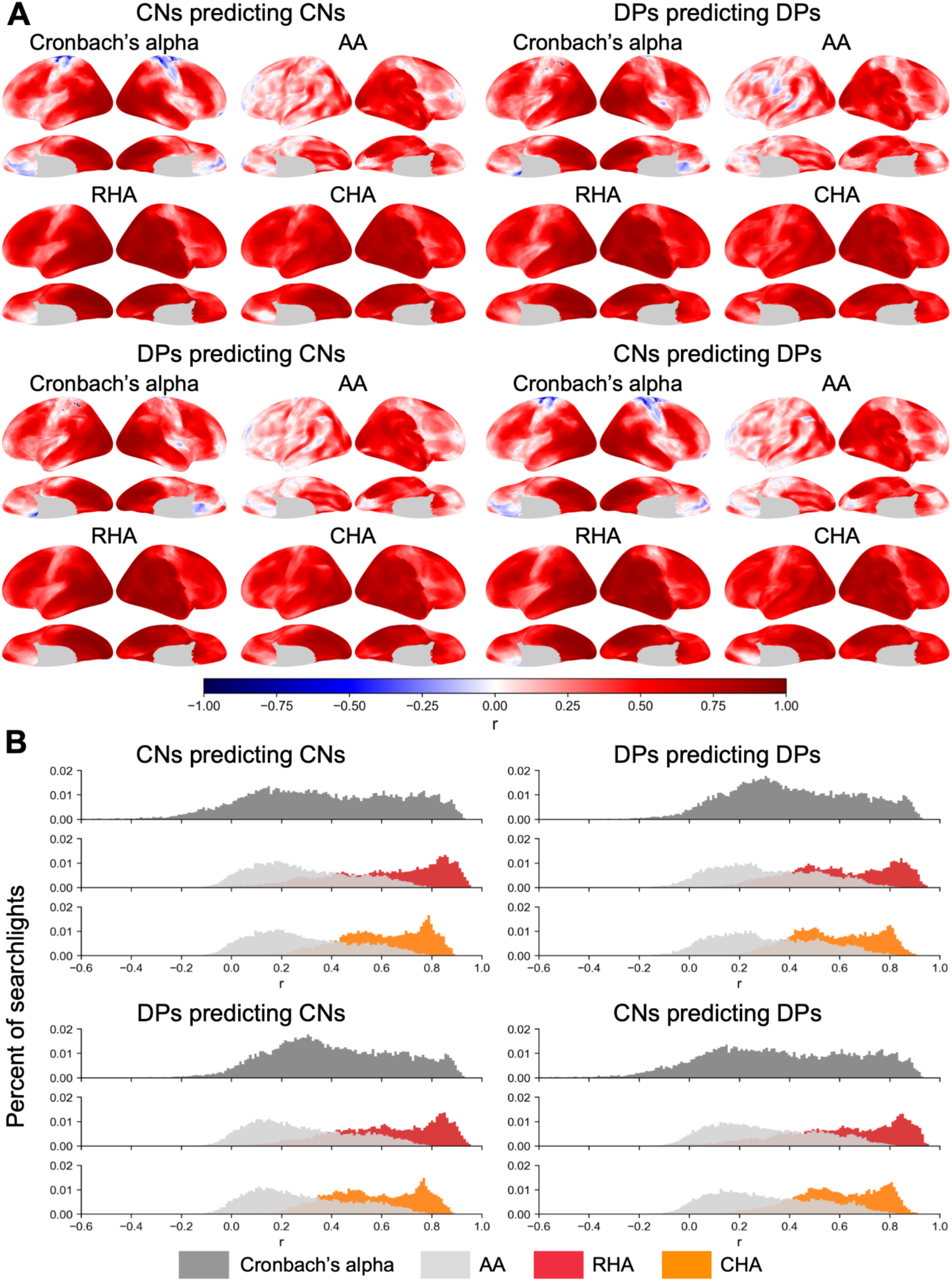
Searchlight analysis of Cronbach’s alpha of face-selective maps and correlations between face-selective maps estimated from participants’ own localizer data and maps estimated from others’ hyperaligned (RHA and CHA) or anatomically aligned (AA) data in the Dartmouth dataset. **(A)** The first two rows display the results of within-group predictions (e.g., predicting control participants’ face-selective maps using data from other control participants). The last two rows display the results of across-group predictions (e.g., predicting control participants’ maps using DPs). For both within- and across-group predictions, Cronbach’s alpha was calculated based on face-selective maps across the five runs in the source participants. The searchlight radius was 15mm. **(B)** Distributions of searchlight correlations of the Cronbach’s alpha, AA, RHA, and CHA maps in Panel A. Within each subplot, the top row represents the distribution of Cronbach’s alphas (dark gray), and the following rows represent the distributions of correlations for RHA (red) and CHA (orange), respectively, plotted against the distribution of correlations for AA (light gray). Similar to Panel A, the top two subplots correspond to within-group predictions and the bottom two subplots correspond to across-group predictions.

To investigate whether an independent task can be used to estimate individualized functional topographies in DPs, we analyzed data from an attentional modulation task (see Methods for design details) scanned with the same group of participants, together with their localizer scans. For each pair of participants, transformation matrices were derived using their attentional modulation task scans and were applied to the localizer runs of the source participant. Similar to the analysis above, the predicted localizer runs were fit with a GLM, and the resulting contrast maps were averaged across all source predictions to obtain the estimated functional topography for the target participant. Because the attentional modulation task scan was not time-locked across participants, it was not appropriate to use RHA. Thus, we only used CHA and AA in this analysis. Our results showed that, with an independent task, CHA significantly outperformed AA in predicting participants’ individualized topographies in both within- and between-group predictions across all categories (*p* < 0.001, Bonferroni-corrected; Supplementary Figures S1 & 2). The predicted topographies with CHA recovered a noticeable amount of individualized features, unlike the blurry topographies predicted with AA (Supplementary Figure S1). Searchlight analysis revealed that areas involved in processing information of the preferred category, including LOC, VT, and STS, demonstrated stronger performance (Supplementary Figure S4), similar to the previous analysis.

Overall, using the Dartmouth dataset, we demonstrated that individualized functional topographies across categories can be estimated with high fidelity from independent functional scans and from other participants in DPs.

### Individualized, high-fidelity predictions using naturalistic stimuli

To evaluate whether high-fidelity predictions of individualized topographies can be achieved with naturalistic stimuli in DPs, we extended our previous analyses to an independent, publicly available dataset with a naturalistic movie scan in DPs (GoT dataset) using similar procedures. This dataset also included one run of a static localizer (faces, scenes, scrambled faces). Following the same hyperalignment pipeline as before, we first derived participant-specific transformation matrices for each pair of participants using the naturalistic movie scan and then applied these transformations to the localizer data of the source participants. Because this dataset contained only a single localizer run, we could not estimate a noise ceiling using Cronbach’s alpha, as this method measures reliability across independent runs. Instead, we computed an inter-subject correlation (ISC) measure by correlating each participant’s contrast map with the other participants’ in a pairwise fashion and averaging the pairwise correlations. This measure estimates between-subject consistency and typically underestimates the noise ceiling. But due to the dataset’s limitations, it provides an alternative solution. In addition, previous studies have demonstrated that dynamic category-selective localizers activate broader regions in the STS and anterior cortices, whereas static localizers more reliably activate posterior areas (Haxby & Gobbini, 2011; Duchaine & Yovel, 2015). As reliable activations were only observed in occipito-temporal regions for the faces-vs-all and scenes-vs-all contrasts, we created an occipital-temporal (OT) anatomical mask encompassing occipital, temporal, and adjacent medial regions (constructed from the Desikan–Killiany Atlas; see Supplementary Figure S6) to focus our evaluation on regions with reliable category-selective responses and performed our correlation analysis within these areas.

Using this dataset with a naturalistic movie scan, we found that hyperalignment successfully recovered individualized topographical information in both DP and control participants, particularly in lateral occipital and ventral temporal areas where activations were most stable across participants (Figure 4). Both RHA and CHA predictions showed significantly higher correlations with participants’ contrast maps estimated from their own localizer run, compared to anatomical alignment (AA), across all contrast types and group comparisons (*p* < 0.001, Bonferroni-corrected; Figure 5; see Supplementary Table 3). The noise ceilings estimated based on pairwise-ISC were well below the other prediction performance values.

**Figure 4.**
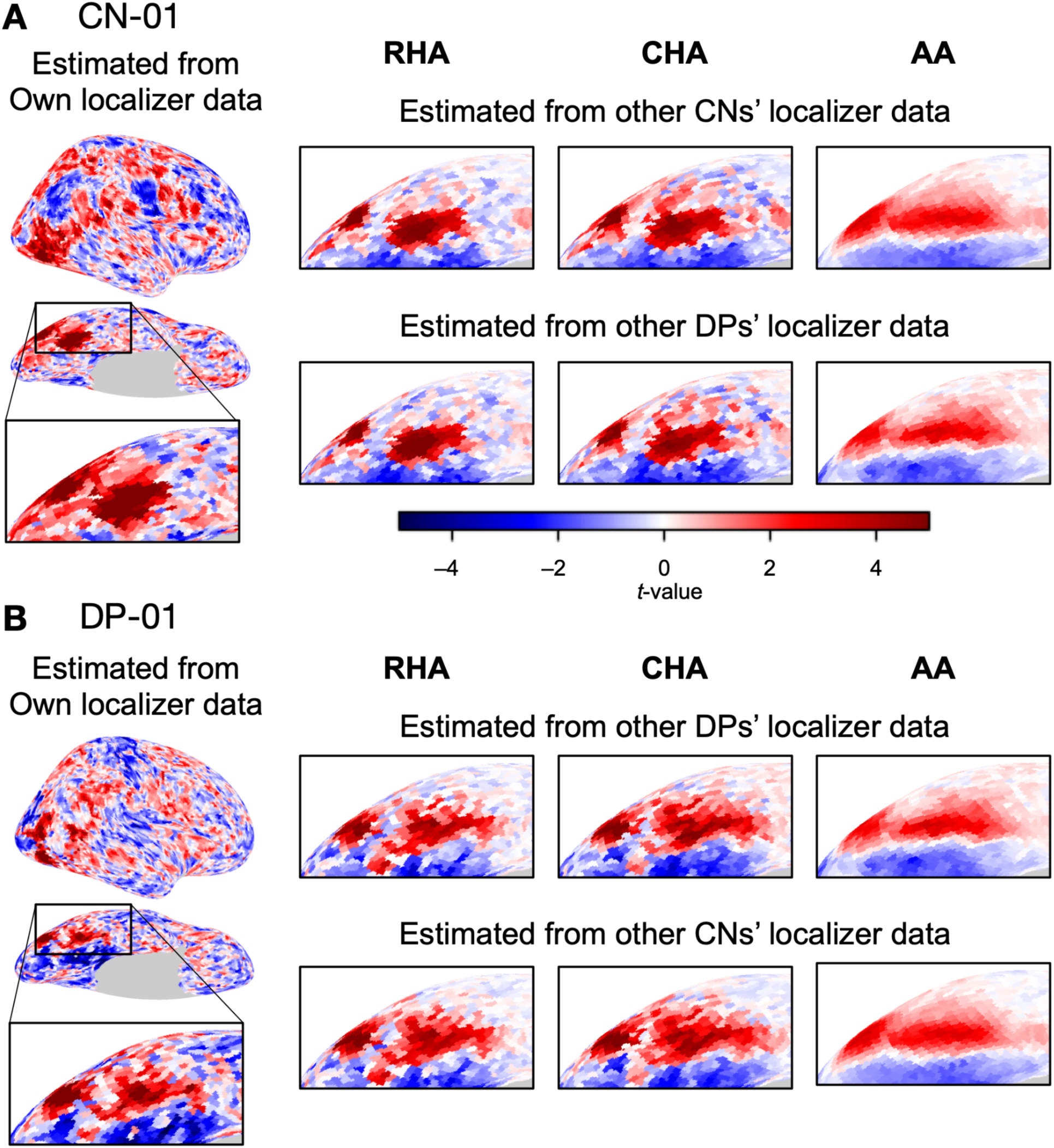
Sample contrast maps and enlarged views of the ventral temporal cortex of control and DP participants in the GoT dataset. Face-selective contrast maps (faces-vs-all) were estimated using the single localizer run in the GoT dataset. **(A)** The estimated face-selective contrast maps for a sample control participant, with expanded views of the right ventral temporal cortex. As shown in Figure 1, the leftmost plot displays the map estimated from that control participant’s own localizer data. The grid shows contrast maps estimated from other participants using either response hyperalignment (RHA), connectivity hyperalignment (CHA), or anatomical alignment (AA). The top row shows maps estimated from other control participants, while the bottom row shows maps estimated from other DP participants. **(B)** The estimated face-selective contrast maps for a DP participant. The left-most plot shows the map estimated from the DP’s own localizer data, while the grid shows maps estimated from other participants. The top row shows estimations from other DP participants, while the bottom row shows estimations from control participants.

**Figure 5.**
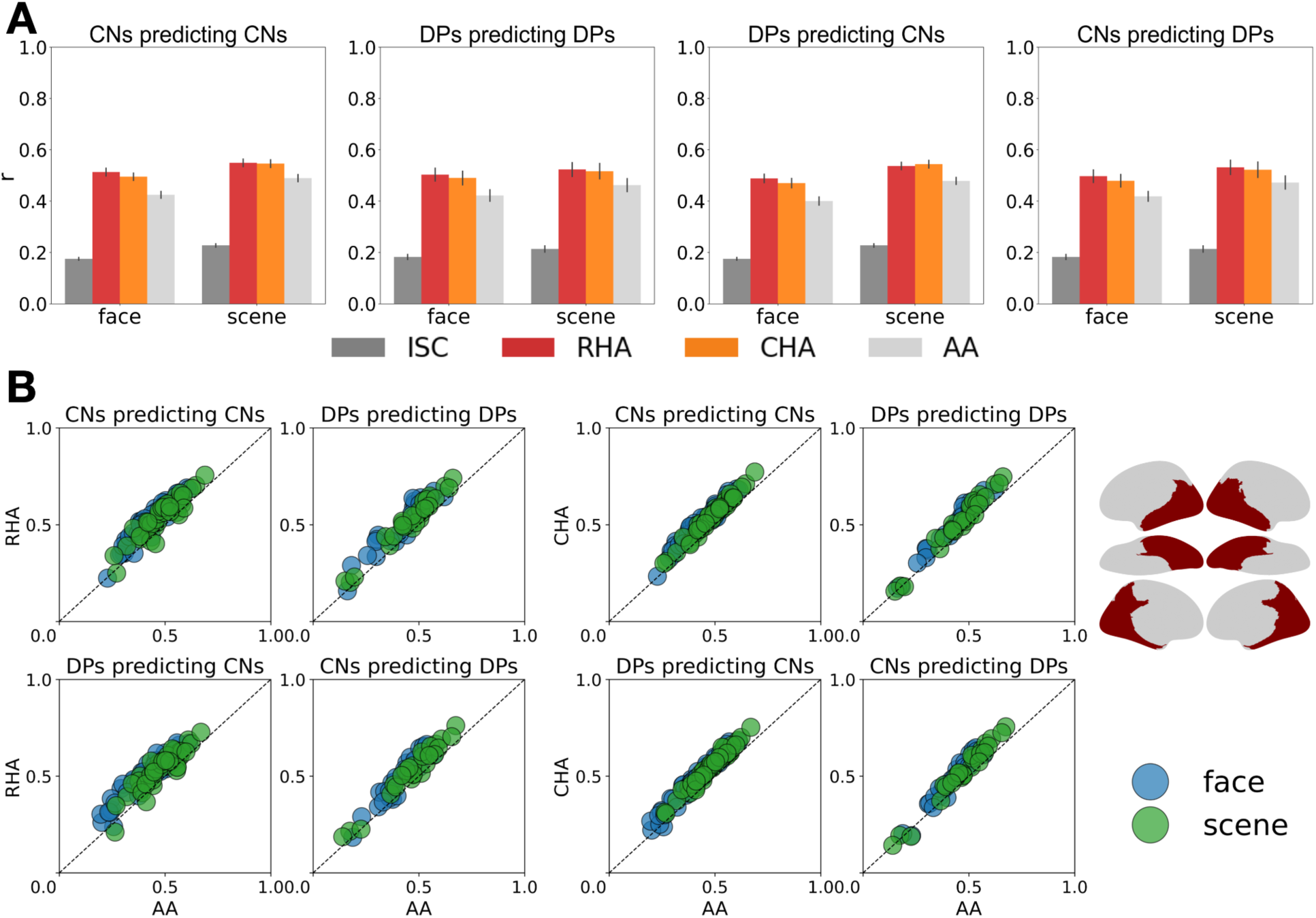
Correlations between contrast maps estimated from participants’ own localizer runs and from other participants’ localizer data across the occipito-temporal cortex (OT) in the GoT dataset. **(A)** Pairwise inter-subject correlation (ISC) values and mean OT correlations for the face and scene categories in the GoT dataset. Each plot shows the average pairwise ISC for both categories (dark gray) and the mean correlations between a participant’s contrast map, estimated from their own localizer runs, and the map estimated from others’ hyperaligned data (RHA and CHA, red and orange, respectively) in OT. Pairwise ISC was calculated using the source group for each prediction. As in Figure 2, the first two plots show within-group comparisons and the last two plots show across-group comparisons. In all plots, hyperalignment predicted results (RHA and CHA) are significantly better than anatomical predicted results (AA, light gray) (p< 0.001, Bonferroni corrected). Error bars stand for ±1 SE. **(B)** Scatter plots of RHA or CHA predicted performance versus AA predicted performance for each individual in OT. Each dot represents a participant’s correlation coefficient between maps estimated using their own localizer data and from others’ using RHA or CHA (y-axis), plotted relative to the correlation coefficient using AA (x-axis). The two left-most scatter plots correspond to RHA, while the two right-most plots correspond to CHA. The brain plot shows the OT mask used in the analysis, and was constructed using anatomical regions from the Desikan-Killany parcellation (aparc.a2009s) to broadly cover the occipital and temporal areas (see also Supplementary Figure S6).

Similarly, we conducted a searchlight analysis to present areas with superior prediction performance. Due to the noise in areas beyond the occipital and ventral temporal cortex in the single-run static localizer data, most noise-ceiling estimate values based on pairwise-ISC were close to zero (Figure 6A, ISC), indicating heterogeneous selective responses across individuals in these areas. Nevertheless, AA correlations were highest in the right hemisphere, and with clusters in bilateral ventral temporal regions (Figure 6A, AA). Compared to AA, both RHA and CHA exhibited improved performance in bilateral occipital and temporal cortices, with peak correlations concentrated in ventral temporal and lateral occipital regions (Figure 6A, RHA, CHA). The corresponding correlation distributions (Figure 6B), restricted to the occipito-temporal mask, revealed clear increases in prediction performance for RHA and CHA, consistent with previous findings. For the scene category, we also found improved performance in OT for RHA and CHA, similar to the face category (Supplementary Figure S5).

**Figure 6.**
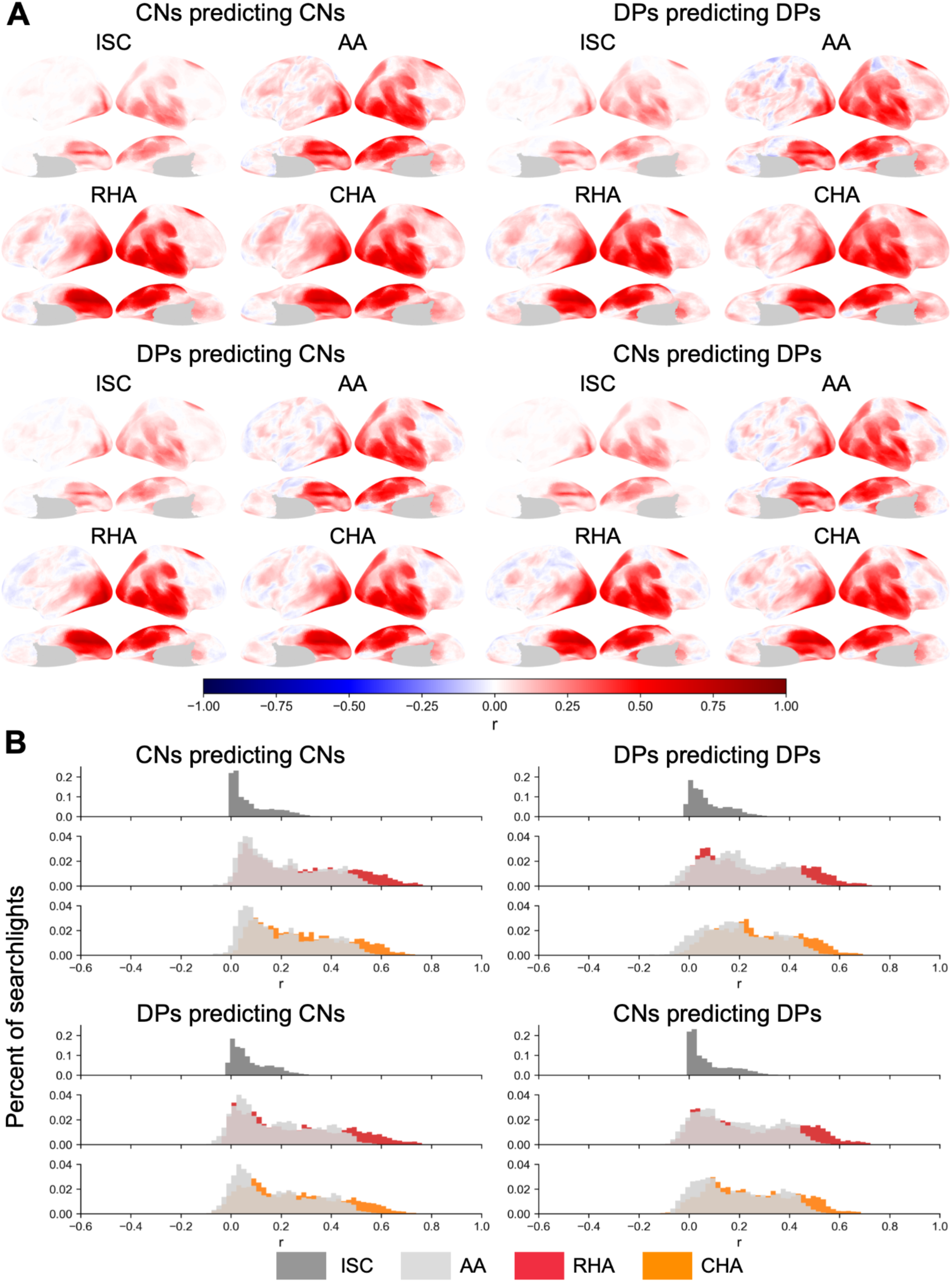
Searchlight analysis of pairwise ISCs of face-selective maps and correlations between face-selective maps from participants’ own localizer data and maps estimated from others’ hyperaligned (RHA and CHA) or anatomically aligned (AA) data in the GoT dataset. **(A)** As in Figure 3, the first two rows display the results of within-group predictions, and the last two rows display the results of across-group predictions. The searchlight pairwise ISC map was calculated by pairwise correlations between participants in the source group for every searchlight, and then averaged across all pairs. The searchlight radius was 15 mm. **(B)** Distributions of searchlight correlations of the pairwise ISC, AA, RHA, and CHA maps in Panel A. Within each subplot, the top row represents the distribution of ISCs (dark gray), and the following rows represent the distributions of correlations for RHA (red) and CHA (orange), respectively, plotted against the distribution of correlations for AA (light gray). Similar to Panel A, the top two subplots correspond to within-group predictions and the bottom two subplots correspond to across-group predictions. The plotted distributions were limited to the occipito-temporal (OT) mask.

### Predicted topographies preserve face-selectivity deficits in DPs

Previous studies have shown that individuals with DP exhibit reduced face-selective activity across the face network, involving regions in the ventral temporal cortex, STS, and frontal lobe (Jiahui et al., 2018; Manippa et al., 2023). The analysis above demonstrated that the framework based on hyperalignment was able to predict cortical topographies in DPs with high fidelity, and we next explored whether these deficits were still preserved using predicted topographies. We used the variable-window method (Norman-Haignere et al., 2016, Jiahui et al., 2018), which defines ROI for each individual by selecting the top X% (5-35% in 5% increments) most category-selective voxels within each anatomical mask drawn at the expected cortex location in a generous manner (Figure 7C, Jiahui et al., 2018). This method was implemented in a leave-one-out fashion. In the Dartmouth dataset, contrast maps from four of the five predicted localizer runs were used to define the most face-selective voxels, and beta weights were extracted from the left-out run in the 12 bilateral ROIs across the face network (Figure 7). For the GoT dataset, which contained only a single localizer run, only the bilateral OFA and FFA were included (Figure 7), and both the most face-selective voxels and the corresponding beta weights were extracted from that run. To make fair comparisons for both datasets, we used CHA-predicted maps.

**Figure 7.**
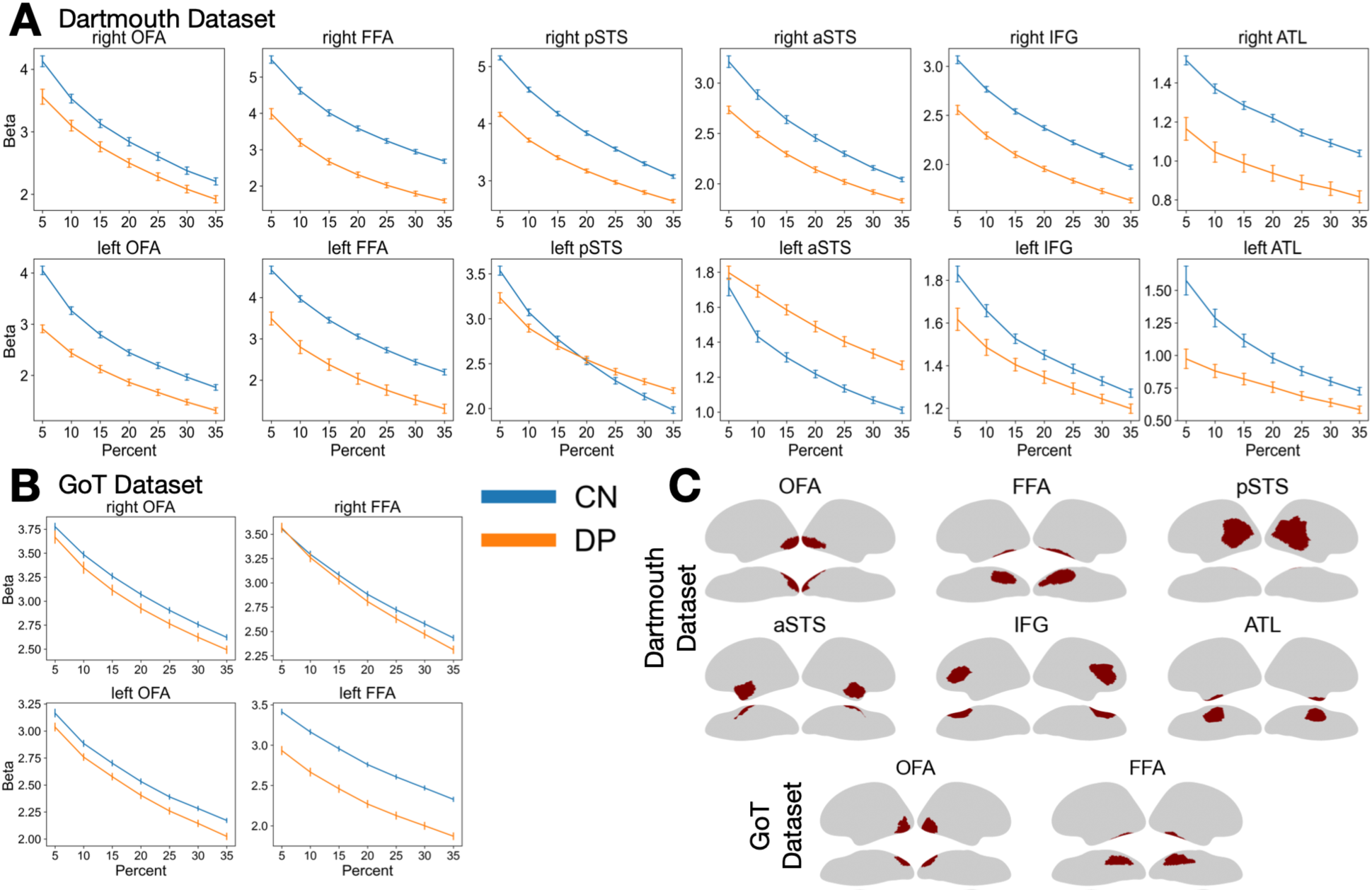
Comparisons of face selectivity between controls and DPs in the face processing network. Face selectivity here refers to the beta weights calculated in the contrast of face vs. all other categories. Beta weights were averaged across vertices and participants within each ROI. **(A)** Face selectivity differences in the Dartmouth dataset using localizer data predicted with CHA from other participants within the same group (Controls-Controls, DP-DP) at ROI sizes from 5 to 35%. For each plot, the mean beta weight (y-axis) was calculated in a cross-validation procedure, by averaging four runs of the face vs. all contrast t-maps, selecting the top X% (5-35%) of voxels (x-axis) within a given anatomical mask (Panel C), and averaging the beta weights within those top voxels in the left-out run in each of the five folds. **(B)** Group-level selectivity differences in the GoT dataset using the predicted localizer runs with CHA. Transformation matrices were derived with the GoT movie scan from other participants in the same group, and the face selectivities were calculated at ROI sizes from 5 to 35%. Two ventral ROIs were reliably localized in this dataset. Since there was only one run of localizer, the average beta weight was calculated by selecting the top X percent of voxels using that run’s face-vs-all t-map within a given anatomical mask (Panel C) and extracting the corresponding beta weights. For both datasets, error bars indicate ±1 SE. **(C)** Anatomical masks for each dataset. The top two rows correspond to masks in the Dartmouth dataset, and the bottom row corresponds to masks in the GoT dataset.

Across all ROIs and sizes in the Dartmouth dataset, DPs showed consistently reduced face-selective responses compared to controls (Figure 7A). These reductions were even more pronounced than those reported in the previous study (Jiahui et al., 2018), likely due to averaging across other participants during the prediction steps. Topographies predicted from participants across groups showed similar though smaller effects (see Supplementary Figure S7A for the across-group analysis). A similar pattern was observed in the GoT dataset as well (Figure 7B & Supplementary Figure S7B). Likewise, these group differences were not as apparent in the original, anatomically aligned topographies (see Supplementary Figure S8).

## Discussion

In this study, we used two datasets with DP participants to estimate individualized category-selective functional topographies using hyperalignment with either task-based scans (Dartmouth dataset; Jiahui et al., 2018, Jiahui et al., 2020) or a naturalistic viewing paradigm (GoT dataset, ds004848; Noad et al., 2024) from other participants. We used the 1-step hyperalignment method (Jiahui et al., 2020; Jiahui et al., 2023) that hyperaligns each source participant to the target participant’s space to derive the transformation matrices, and applied the transformations to the localizer data of the source participant. We found that, similar to previous work in typical participants, individualized functional topographies were estimated in high fidelity in DPs using hyperalignment across face-, body-, scene-, and object-selective topographies. Whole-brain correlation and searchlight analyses found that these estimations were highly correlated with individuals’ topographies estimated from their own localizer data, especially in areas known to show strong category-selective responses. We also found that predictions using out-of-group participants, such as predicting DPs’ topographies from control participants and vice versa, provided comparable prediction performances to within-group estimations. Notably, we found that the estimated topographies also preserved the face selectivity reductions in DPs’ face-selective network that have been previously reported in the literature (see Manippa et al., 2023 for a review). Our findings support a framework for individualized estimation of category-selective functional topographies in DPs with naturalistic or task scans using hyperalignment without a separate independent localizer scan, and lay the foundation for future generalizations to other neuropsychological and clinical groups.

Localizing individualized functional areas has been a central problem in the history of modern cognitive neuroscience. Functional localizers, which are usually a separate task with block-designed scans, are performed as the classic method to achieve this goal (Saxe et al., 2006; Fedorenko et al., 2011). Though there has been a growing trend to include more stimulus categories and cognitive functions within a single localizer scan (Marvi et al., 2025, Moreno et al., 2025), most of these localizers are typically targeted at one type of cognitive function, adding a significant amount of time to the scanning schedule, especially when multiple types of functions are preferred. Additionally, localizer scans are best designed for healthy young adults, making individualized localization challenging in other populations, such as clinical and neuropsychological groups, and across the age spectrum, such as children. Previous research into typical participants has shown that it is possible to estimate participants’ functional topography based on a non-localizer task (Jiahui et al., 2020; Jiahui et al., 2023). For example, if a previously-collected dataset already contains functional localizer scans and a naturalistic movie, functional localizer data can be estimated for a new set of participants using transformation matrices between the old task and new task scans with hyperalignment. For DP, this presents a promising avenue for research. DPs show impaired performance in face recognition, and deficits in face-selectivity (Avidan & Behrmann, 2009; Furl et al., 2011; Dinkelacker et al., 2011;, Avidan et al., 2014; Jiahui et al., 2018), connectivity (Gomez et al., 2015; Song et al., 2015; Lohse et al., 2016; Rosenthal et al., 2017; Zhao et al., 2018), and multi-voxel patterns (Rivolta et al., 2014; Zhang et al., 2015; Tian et al., 2020; Haeger et al., 2021; Zhao et al., 2022) have been observed in the face-selective network. Beyond faces, DPs sometimes also showed atypical performance in body perception (Righart & de Gelder, 2007; Biotti et al., 2017; Rivolta et al., 2017), spatial navigation and topographic memory (Duchaine & Nakayama, 2005; Klargaard et al., 2016), and some object recognition (Duchaine & Nakayama, 2005; Behrmann et al., 2005; Duchaine & Nakayama, 2007; Geskin & Behrmann, 2020). These avenues have been relatively less explored compared to face perception and face selectivity. In addition to high-level visual categories, few studies have investigated whether DPs are impaired in other cognitive functions such as theory of mind, speech processing, and calculation. To properly examine the underlying neural mechanisms of these different cognitive abilities, using individualized localizations with traditional functional localizers will significantly increase scanning time, making feasibility even more difficult. Our new framework enables the estimation of various individualized functional topographies in DPs using existing datasets with localizer scans. This paves the way for future DP research to identify different types of individualized areas from a separate scan, such as a movie. This framework also enables the integration of existing DP datasets across studies, allowing the existing small-sample datasets from atypical populations to be pooled into larger cohorts to improve the scalability of research in these groups.

Interestingly, we observed similar cross-group prediction results compared to within-group predictions. This result suggests that it is possible to estimate DP’s individualized topographies from a group of typical participants. Previous work has shown that the general organization of the category-selective areas was normal and similar to that of typical participants (Jiahui et al., 2018). The successful cross-group predictions further demonstrate the preserved general organizations in this group, given their severe deficits in face recognition. It is worth noting that the general organization of category-selective topographies is remarkably stable across typical individuals (Peelen & Downing, 2017; Rosenke et al., 2021) as well as in many different types of psychiatric disorders, such as autism (Hadjikhani et al., 2004; Nickl-Jockschat et al., 2015; Kliemann et al., 2018; Vandewouw et al., 2020; D’Mello et al., 2023) and schizophrenia (Yoon et al., 2006; Bortolon et al., 2015; Kovács et al., 2019; Lee et al., 2019). However, our ROI analyses comparing face selectivity between the typical and DP groups showed that using predictions derived from the control group for DPs reduced the group-level differences, such that the predictions shifted DP selectivity towards the control profile (Supplementary Figure S7). Hence, while we have shown that it is possible to predict DP topographies using typical participants, extra considerations should be taken when conducting investigations into category selectivity beyond topographical organization. Ideally, a normative dataset of DPs (or other neuropsychological or clinical groups in the general case) should be used to derive predictions in order to preserve characteristics beyond the general architectures and conduct scalable investigations of underlying neural mechanisms.

Similar to previous studies, we found significant differences between face-selective responses using dynamic and static localizers. In the Dartmouth dataset, dynamic stimuli elicited reliable selectivity in both ventral and dorsal face areas, as well as the frontal areas. In the GoT dataset, the static localizer only reliably captured FFA and OFA. These findings align with previous research that the ventral pathway in the face-selective network is more involved in processing static features of faces, and the dorsal pathway is more important for the dynamic aspects of faces (Haxby and Gobbini, 2011; Duchaine & Yovel, 2015; Bernstein et al., 2018; Pitcher et al., 2019). Previous studies have demonstrated alterations in functional connectivity and face selectivity in the anterior areas (Avidan & Behrmann, 2009; Dinkelacker et al., 2011; Avidan et al., 2014, Lohse et al., 2016, Rosenthal et al., 2017, Jiahui et al., 2018), yet the functional contributions of these areas are still not fully explored. For example, Avidan & Behrmann (2014) have hypothesized that the face-selective areas in the prefrontal cortex, which show deficits in DP, may be involved in holistic/configural face processing, as a TMS study reported a double dissociation between right and left dorsolateral prefrontal cortex and configural versus featural processing (Renzi et al., 2013). This area may also be implicated in the processing of eye-gaze information (Chan & Downing, 2011) and identity (Guntapalli et al., 2017). Hence, reliably localizing individualized face areas, especially the anterior areas, will be important in future research, and dynamic localizers will be a superior alternative to static ones.

We were able to successfully predict category-selective topographies using both naturalistic viewing movie scan and a task-based scan. As demonstrated in previous studies, hyperalignment works best with naturalistic stimuli, as naturalistic stimuli contain rich information beyond controlled experiments that engage multiple functional systems at the same time (Haxby et al., 2011; Guntapalli et al., 2016). In addition, naturalistic stimuli often generate higher-quality data (e.g., reduced head motion, better attention from participants) than controlled experiments across different participant groups (Hasson et al., 2004, 2010; Vanderwal et al., 2015, 2017, 2019; Redcay & Moraczewski, 2020; Haxby et al., 2020, Nastase et al., 2020; Finn, 2021; Kamps et al., 2022). While task-based scans can still be used in the prediction with hyperalignment, their efficacy and generalizability are constrained by the design and the scope of the task (Haxby et al., 2020). Algorithm-wise, RHA requires participants’ response time series to be time-locked, which is not the case in most of the tasks, and only CHA can be used. For example, in the Dartmouth dataset, although the block order of the attentional modulation task was identical across individuals, the stimuli within each block were randomized. In principle, such stimulus-level randomization in a block design is typically a minor issue and would not substantially affect time-locked analyses. However, more importantly, DP participants were instructed to attend to the same facial dimension (e.g., face identity) as typical participants, but due to their deficits in face recognition, they may have had difficulty following the task instructions. As a result, their neural responses during this task were likely more idiosyncratic compared to those of typical participants. Under these circumstances, it is not appropriate to assume time-locked response time series across participants for this task. In addition to rich information and better data quality, naturalistic viewing paradigms typically do not suffer from such a constraint, since all participants watch the same movie. It is worth noting that commercial movies with continuous narratives often better align individuals and can lead to stronger cross-individual predictions than short clips concatenated together (Hasson et al., 2008; Ngyuen et al., 2019). This may represent an additional factor contributing to the weaker prediction results in the GoT dataset, beyond the limitation of having only a single localizer run, as the movie stimuli in the GoT dataset consist of clips drawn from multiple segments of the TV series.

It has been a frequently asked question about the length of the movie to ensure proper estimation of individualized topography. In previous work, full-length movies or half-length movies with about one hour of scanning time were used to derive transformation matrices with hyperalignment, and they generated robust estimations of retinotopic and category-selective maps (Guntapalli et al., 2016; Jiahui et al., 2020; Jiahui et al., 2023; Feilong et al., 2023). While there is no established minimum scan time, systematic analyses suggest that longer durations generally improve estimation reliability. Nevertheless, approximately 10-15 minutes of movie viewing is often sufficient (Jiahui et al., 2020; Feilong et al., 2023). In this analysis, successful predictions were obtained using the approximately 13-minute movie from the GoT dataset, which consists of concatenated segments, demonstrating that reliable predictions can still be achieved with relatively short naturalistic movie scans.

Future work can extend this framework in several directions. Because cross-group hyperalignment produced robust predictions and allows category-selective topographies to be estimated from scans not originally designed as functional localizers, existing datasets from typical participants and other tasks can help estimate individualized topographies in small atypical datasets, enabling currently existing DP datasets to be more meaningfully integrated and analyzed to obtain more robust findings. However, to properly characterize population-specific neural features such as reduced face selectivity in DP, larger normative datasets built specifically for the target population will be important. Naturalistic paradigms will be particularly useful in this context because they engage multiple functional systems simultaneously and are much more friendly for clinical or neuropsychological populations. In summary, our results demonstrate that hyperalignment can estimate individualized category-selective functional topographies in high fidelity in DPs using either task-based scans or naturalistic movie viewing, providing a scalable framework for studying the neural basis of DP and other atypical populations.

## Methods

### Participants

#### Dartmouth Dataset

We used a previously-collected dataset from Dartmouth College (Dartmouth Dataset, Jiahui et al. 2018, Jiahui et al, 2020), which consisted of twenty-two DPs (seven males, mean age 41.9 y) and 25 typical adults (10 males, mean age 42.3 y). The DPs were recruited from www.faceblind.org, and their face recognition abilities were assessed with the Cambridge Face Memory Test (CMFT) (Duchaine and Nakayama, 2006), a famous face test (Duchaine and Nakayama, 2005), and an old-new face discrimination test (Duchaine and Nakayama, 2005) One DP performed slightly above the diagnostic cutoff line (2 or more SDs below the mean of controls in at least two of the three tests), but performed poorly (CFMT: z = −1.9; famous face: z = −7.1; old-new: z =−0.5), and was also included. All participants had normal or corrected-to-normal vision and had no current psychiatric disorders when data were collected. Participants provided written informed consent before doing the tasks, and all procedures were approved by Dartmouth’s Committee for the Protection of Human Participants.

#### Game of Thrones (GoT) Dataset

We also used a publicly available dataset from OpenNeuro, the Game of Thrones Dataset (ds004848, Noad et al., 2024). This dataset consisted of twenty-eight DPs (12 males, median age 47 years) and 45 typical adults (15 males, median age 19 years). DP participants were recruited through www.troublewithfaces.org and other online sources, and were diagnosed with the PI20, a 20-item index to measure self-reported face recognition abilities, and the CMFT. Participants who scored above 65 on the PI20 and below 65% on the CMFT were considered prosopagnosic (see Noad et al., 2024 for details).

### fMRI scanning

#### Dartmouth Dataset

There were two sets of scans in the Dartmouth Dataset. The first scan consisted of a dynamic localizer, which had five categories: faces, scenes, bodies, objects, and scrambled objects. Faces and objects stimuli were selected from stimuli used in Fox et al. (2009), scene and body video clips were from Pitcher et al. (2011), and scrambled objects were created by spatially scrambling the video clips of the objects into 24 × 16 grids (Jiahui et al., 2018). There were five runs in the localizer, each lasting 4.2 minutes. Each run consisted of ten 12s stimulus blocks interleaved by 12s fixation blocks, and each category was displayed twice in a quasi-random order across scans. In each stimulus block, six 1.5s video clips were presented, interleaved by 500ms blank fixation screens. Participants were asked to complete a one-back task during the scan to maintain their attention.

The second set of scans was an attentional modulation task. The task was also block-designed and included six runs in total. In each run, nine stimulus blocks that lasted 18s each were interleaved with 18 s fixation blocks. The experiment included three conditions (identity, expression, and view). Before each stimulus block began, a word that indicated the condition of the block was presented in the center of the screen for three seconds. Participants pressed a button when the target aspect of a block (identity, expression, or view) was the same across two trials in a row (Jiahui et al., 2020).

Participants were scanned using a 3.0-T Phillips MR scanner (Philips Medical Systems, WA, USA) with a SENSE (SENSitivity Encoding) 32-channel head coil (see Jiahui et al., 2018 for more details). An anatomical volume was acquired using a high-resolution 3D magnetization-prepared rapid gradient-echo sequence (220 slices, field of view = 240 mm, acquisition matrix = 256 × 256, voxel size = 1 × 0.94 × 0.94 mm) at the beginning of the scan. Functional images were collected using echo-planar functional sequence (TR = 2000ms, TE = 35 ms, flip angle = 90°, voxel size = 3 × 3 × 3 mm) (See Jiahui et al., 2018 for more details).

#### Game of Thrones (GoT) Dataset

Two sets of scans were also in the Game of Thrones dataset. The first scan consisted of a static localizer with three categories: faces, scenes, and phase-scrambled faces. Face stimuli were taken from the Radboud database (Langner et al., 2010), while scene stimuli consisted of indoor, outdoor man-made, and outdoor natural stimuli from the SUN database (Xiao et al., 2010). There was one run of the localizer, which consisted of nine blocks in a pseudorandomized order. Each block contained four images of a category that were presented for 600 ms with a 200 ms inter-stimulus interval (ISI), for a total of nine seconds per block. The total scan time was 244 seconds. To maintain attention, participants responded to periodic color changes in the fixation cross via button press.

The second scan was a naturalistic viewing paradigm where participants watched a movie made up of clips from the television show Game of Thrones. This was one run of passive movie viewing made up of ten distinct clips ranging from 50 to 117 s, for a total run time of 778 s (12 m 58 s). Participants were also tested on their familiarity with the Game of Thrones show after the scan based on character, scene, and narrative knowledge, which resulted in two sets of groups in controls and DPs that were either familiar or unfamiliar with GoT.

All scans were conducted at the York Neuroimaging Centre using a 3 T Siemens Magnetom Prisma MRI scanner equipped with a 64-channel phased array head coil. Data were collected via a gradient-echo echoplanar imaging (EPI) sequence from 60 axial slices, with parameters: TR = 2 s, TE = 30 ms, FOV = 240 × 240 mm, matrix size = 80 × 80, voxel size = 3 × 3 × 3 mm, slice thickness = 3 mm, flip angle = 80°, phase encoding from anterior to posterior, and a multiband acceleration factor of 2. T1-weighted structural images were obtained from 176 sagittal slices (TR = 2,300 ms, TE = 2.26 ms, matrix size = 256 × 256, voxel size = 1 × 1 × 1 mm, slice thickness = 1 mm, flip angle = 8°). Additionally, field maps were collected from 60 slices (TR = 554 ms, short TE = 4.90 ms, long TE = 7.38 ms, matrix size = 80 × 80, voxel size = 3 × 3 × 3 mm, slice thickness = 3 mm, flip angle = 60°) (see Noad et al., 2024 for more details).

### MRI preprocessing

We followed standard preprocessing pipelines for both datasets using fMRIPrep version 24.1.1 (Jiahui et al. 2023; Esteban et al., 2019). Volumes were corrected for head motion using rigid-body realignment and coregistered to the corresponding T1-weighted anatomical image. Susceptibility distortions were corrected using fMRIPrep’s fieldmap-less susceptibility distortion correction (SDC) method based on SyN registration (Esteban et al., 2019). Functional data were projected onto the *fsaverage* cortical surface using FreeSurfer’s surface-based registration (Fischl et al., 1999). Data were further resampled to *fsaverage5* space, a cortical mesh with 20,484 vertices in both hemispheres (Jiahui et al., 2023). The functional data were further denoised by regressing out six motion parameters and their temporal derivatives, the global signal, framewise displacement, the first six components from white matter from aCompCor, and polynomial trends up to the second order (Jiahui et al., 2020, 2023, Power et al., 2014, Behzadi et al., 2007). After denoising, the medial wall vertices were removed.

### Searchlight Hyperalignment

#### Connectivity Hyperalignment (CHA)

To conduct searchlight connectivity hyperalignment, we first calculated whole-brain functional connectivity using vertices on a sparser cortical surface as the connectivity targets. In particular, we used *fsaverage3*, which corresponds to 642 vertices per hemisphere before removal of the medial wall. The seeds were the vertices on the original cortical surface in *fsaverage5* space. We averaged the time series in a 13 mm searchlight centered on the connectivity targets and built the connectivity profile by calculating connectivity between the mean time series and the original time series of the connectivity seeds (Jiahui et al., 2020, Jiahui et al., 2023).

To perform CHA, we projected one participant’s connectome into another participant’s space. For each 15 mm searchlight, the transformation matrix was derived by aligning a given source participant’s connectome to a target participant’s using the Procrustes transformation, and transformation matrices were aggregated across all searchlights in each hemisphere to obtain one transformation matrix for each pair of participants (Guntupalli et al., 2016). This transformation matrix was then applied to the localizer runs of the source participant to align the localizer data to the target participant’s space. We did this for all possible combinations of source-target participants (Jiahui et al., 2020, Jiahui et al., 2023). For hyperalignment involving the localizer data only, we split the localizer runs in a leave-one-out fashion, derived transformation matrices using a concatenation of four of the runs and predicted the left-out run. For analysis involving the attentional modulation experiment or GoT movie viewing, we concatenated all six runs of the experiment or used the whole movie scan as is, respectively, to derive the transformation matrices.

#### Response Hyperalignment (RHA)

For RHA analysis, the same steps were used as in CHA. The main difference is that RHA takes the response pattern vectors (i.e. the timeseries) as the input, instead of the connectivity pattern vectors, within a given searchlight. RHA requires that the inputs be time-locked; therefore, we only applied RHA to hyperalign participants using localizer scans in the Dartmouth dataset and to the naturalistic movie-watching data in the GoT dataset.

### General Linear Model

We estimated each participant’s category-selective maps by calculating univariate contrasts using a General Linear Model (GLM) with their own localizer runs and localizer runs estimated from other participants. Each run was analyzed separately, and the results were averaged across all runs. Specifically, contrast maps for each category (faces, bodies, scenes, and objects) were derived by calculating the contrast of the target category vs. all the other categories.

#### ROI Analysis

We followed the pipeline described in Jiahui et al. (2018) to localize functional ROIs, aiming to decrease subjectivity in the traditional threshold-based localization process and to better handle DPs who responded weakly. For the Dartmouth Dataset, each functional mask was manually prepared by referring to the category-selective voxels of both groups at a liberal threshold (p < 0.05) to include as many possible vertices. The individual ROI for each participant was then defined using the “variable-window method” to find the 5-35% most category-selective voxels within the mask (Jiahui et al., 2018). To avoid the double-dipping problem (Kriegeskorte et al., 2009), all analyses were based on a leave-one-run-out process, in which four runs were used for localization and one run was used for extracting beta weights. For the GoT dataset, the same process was performed. However, due to the limitation of the localizer data in the GoT dataset (e.g., static localizer, only one scan run), only FFA, OFA for faces and PPA, RSC for scenes were able to be localized (Supplementary Figure S6), and no splitting runs were performed in the following analysis.

### Whole-brain and searchlight analyses

To evaluate the performance of the prediction, we conducted both whole-brain and searchlight-based correlation analyses. For our whole-brain correlations, we calculated the correlation between a given participant’s contrast map estimated from their own localizer data and the predicted contrast maps from RHA, CHA, or anatomical alignment (AA). We tested both within-group (i.e. controls to controls, DPs to DPs) and across-group (controls to DPs, and vice versa) predictions, since we wanted to examine if it was possible to generate predicted topographies using participants from the other group. We used repeated-measures t-tests to examine the performance of hyperalignment (RHA, CHA) versus AA. We also calculated a noise ceiling for the Dartmouth dataset by using the whole-brain maps from the five runs of a participant’s localizer data to calculate Cronbach’s alpha, and averaged across participants within their respective group. For the GoT dataset we used intersubject correlation (ISC) in lieu of Cronbach’s alpha to calculate a noise ceiling since each subject only had one run of the localizer. Specifically, for a given participant, we calculated the pairwise ISC of their contrast map with every other participant in their group, and averaged across all pairwise correlations to have a single ISC value for that participant. Given that this one-run static localizer was only able to reliably localize regions in the occipital and ventral temporal cortices, we created a bilateral occipito-temporal mask using contiguous regions from the Desikan-Killany parcellation (aparc.a2009s; Destrieux et al., 2010) that broadly covered both face- and scene-activated areas from the localizer data (Supplementary Figure S6).

We also conducted a searchlight analysis to demonstrate the prediction performance across the cortex. Following similar analysis steps, we correlated the contrast map estimated using a given participant’s own localizer data with the maps estimated using RHA, CHA, and AA within each 15mm searchlight. We also calculated Cronbach’s alpha and pairwise ISC within 15mm searchlights for the Dartmouth dataset and GoT dataset, respectively. This was conducted for both within-group and across-group predictions. For all across-group analyses, the noise ceiling measurement was based on participants who were used to calculate the predicted map. For example, for the prediction from controls to DPs, the Cronbach’s alpha values/pairwise ISCs were calculated based on the control group.

## Acknowledgements

This work was supported by a startup grant to G.J. from the University of Texas at Dallas.

## Supplementary Materials

**Figure S1.**
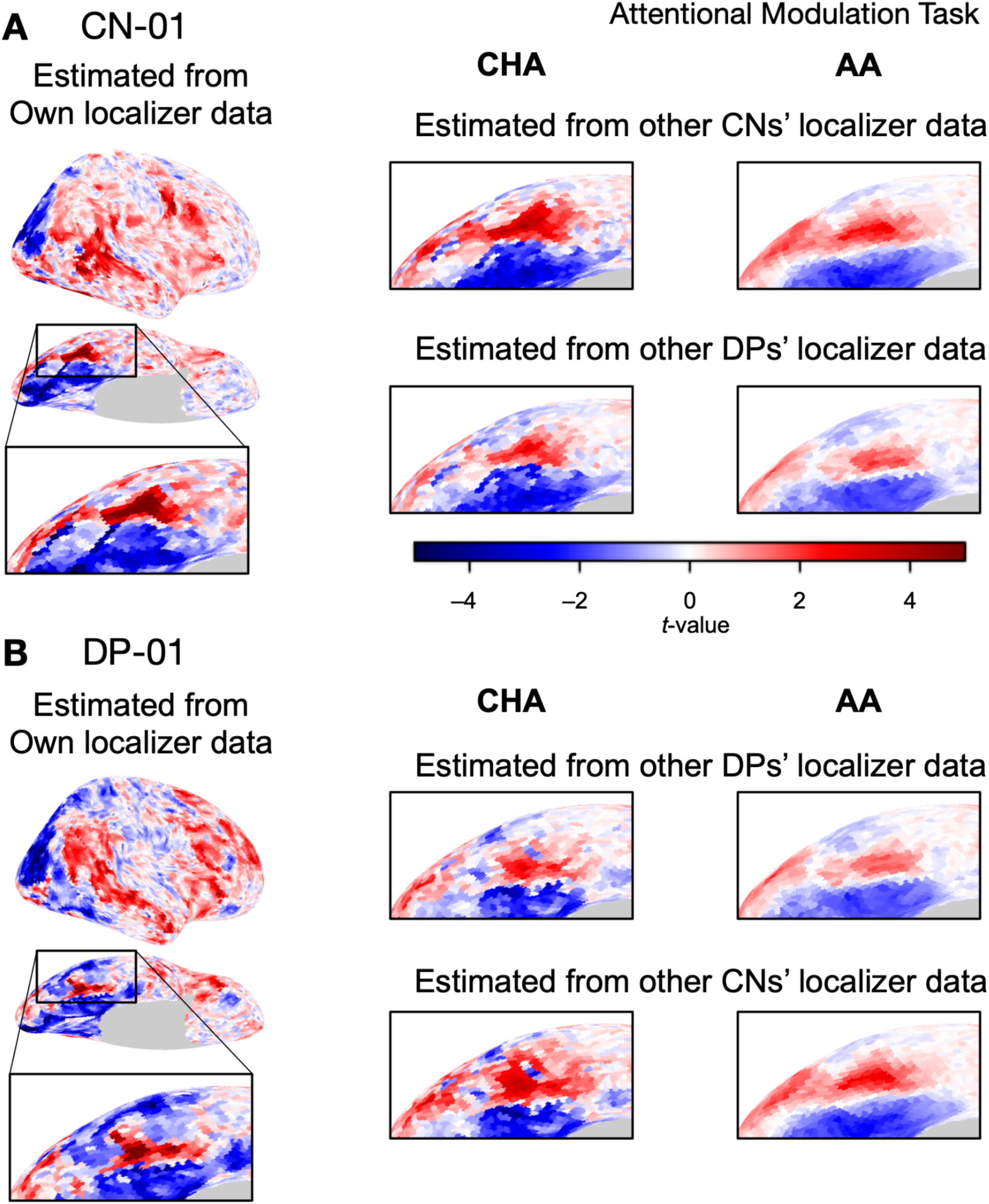
Sample contrast maps and enlarged views of the ventral temporal cortex of control and DP participants. Contrast maps for face-selective topographies (faces-vs-all) were estimated using localizer runs from the Dartmouth dataset. Here we used the attentional modulation scan in the dataset to derive transformation matrices with hyperalignment. **(A)** The estimated face-selective contrast maps for a sample control participant, with expanded views of the right ventral temporal cortex. The left-most plot shows the map estimated from that control participant’s own localizer data. The grid shows contrast maps estimated from other participants using connectivity hyperalignment (CHA) or anatomical alignment (AA). The top row shows maps estimated from other control participants, while the bottom row shows maps estimated from other DP participants. **(B)** The estimated face-selective contrast maps for a sample DP participant. As in **(A)**, the left-most plot shows the map estimated from the DP’s own localizer data, while the grid shows maps estimated from other participants. The top row shows estimations from other DP participants, while the bottom row shows estimations from control participants. RHA was excluded from these plots because the attentional modulation task did not produce time-locked response time series and was therefore not suitable for RHA.

**Figure S2.**
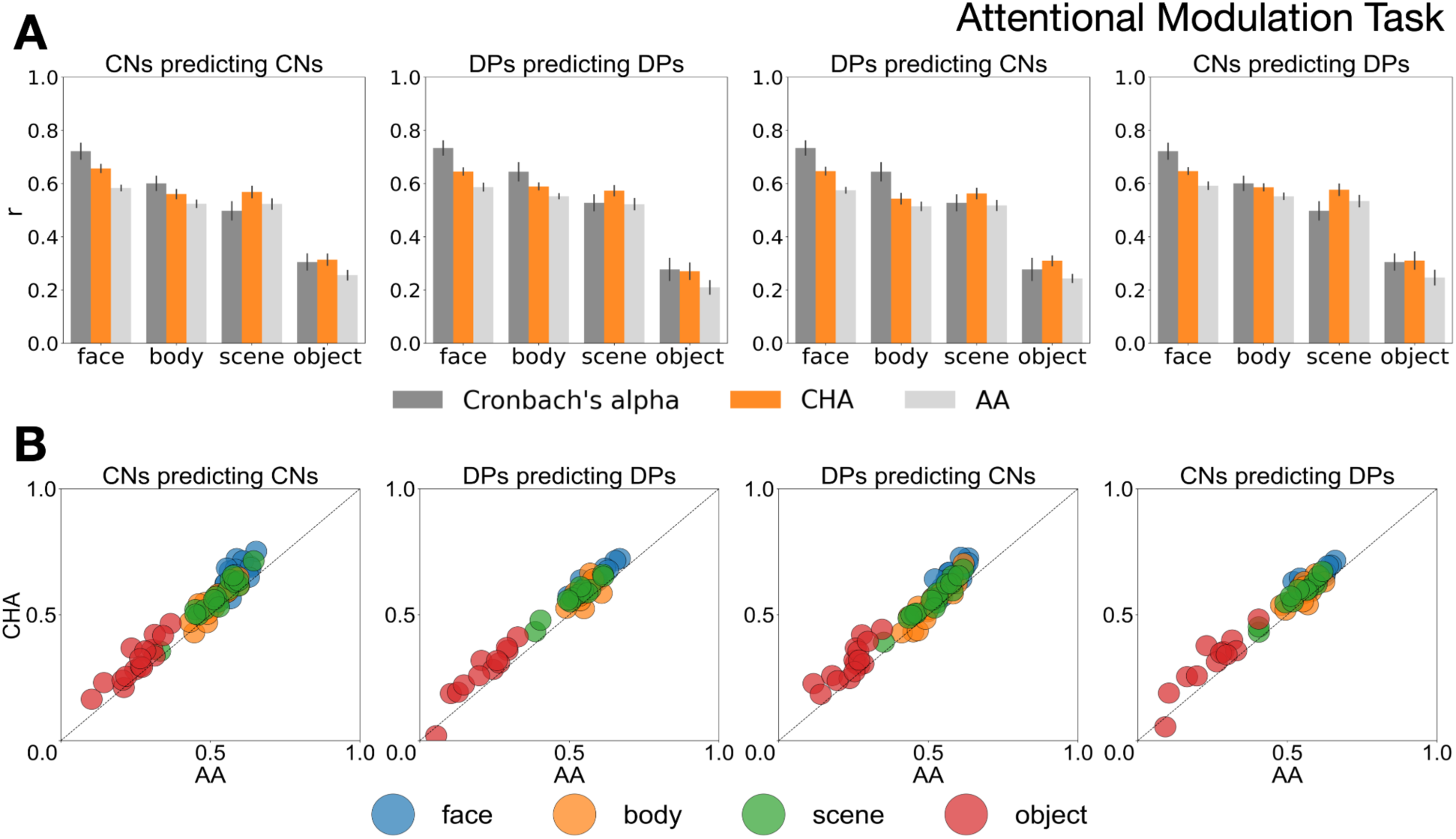
Correlations between contrast maps estimated from participants’ own localizer runs and from other participants’ localizer data. Predicted localizer runs were derived using hyperalignment with the attentional modulation task. **(A)** Cronbach’s alpha values and mean whole-brain correlations across the four categories in the Dartmouth dataset. Each plot shows the Cronbach’s alpha values for each category and mean correlations between a participant’s contrast map and their predicted map from connectivity hyperalignment (CHA) or anatomical alignment (AA). Cronbach’s alpha was calculated based on data from the source group for each prediction. From left to right, the first two plots show within-group comparisons (Controls to Controls, DPs to DPs), while the last two plots show across-group comparisons (DPs to Controls, Controls to DPs). In the across-group comparisons, control participants’ data were hyperaligned to DP participants’ space (and vice versa). Since participants were not time-locked in this task, they could not be aligned using RHA. The CHA predicted results are significantly better than AA predicted results (*p*< .05, Bonferroni corrected). Error bars stand for ±1 SE. **(B)** Scatter plots of CHA predicted performance versus AA predicted performance for each individual. Each dot represents a participant’s correlation coefficient between maps estimated using their own localizer data and using CHA (y-axis), plotted relative to the correlation coefficient using their AA (x-axis).

**Figure S3.**
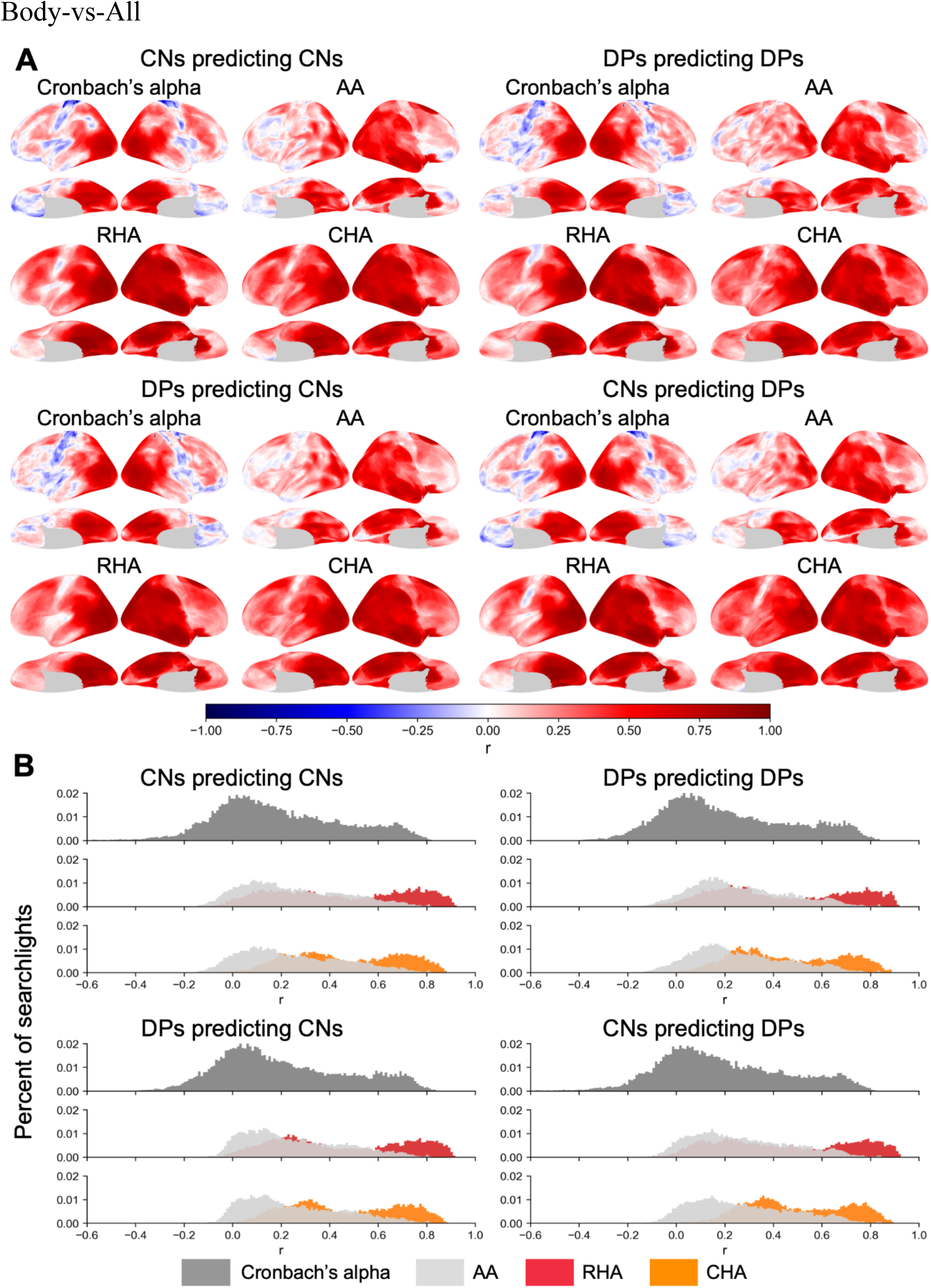
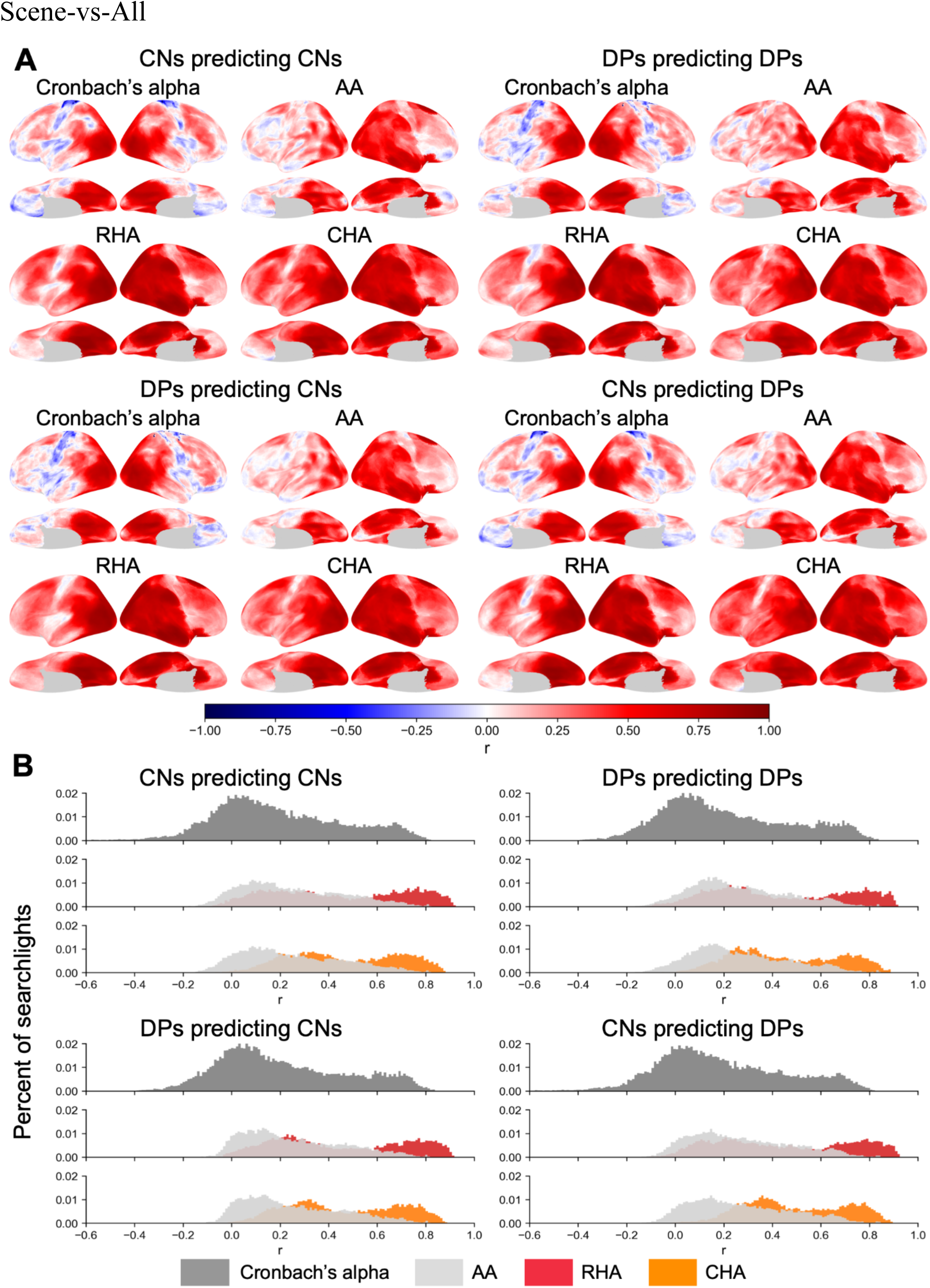
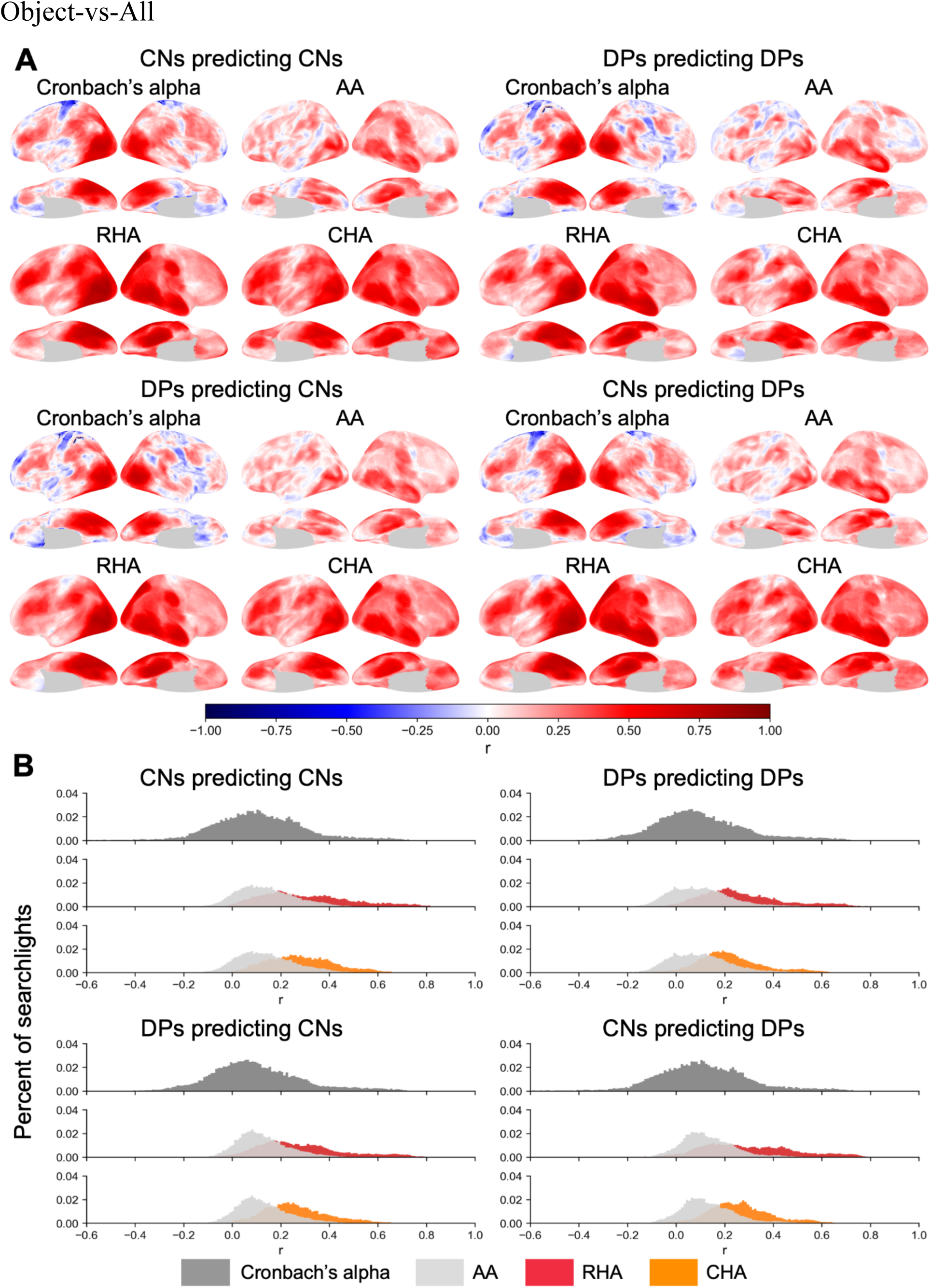
Searchlight analysis of Cronbach’s alpha of body-, scene-, and object-selective maps and correlations between category-selective maps from participants’ own localizer data and maps estimated from others’ hyperaligned (RHA and CHA) or anatomically aligned data in the Dartmouth dataset. For all categories, the contrast was defined as category vs. all other categories and a 15mm searchlight radius was used. **(A)** The first two rows display the results of within-group predictions (e.g. predicting control participants’ face-selective maps using data from other control participants). Cronbach’s alpha was calculated based on category-selective maps across the five runs of control participants. The last two rows display the results of across-group predictions (e.g. predicting control participants’ maps using DPs). **(B)** Distributions of searchlight correlations of the Cronbach’s alpha, AA, RHA, and CHA maps in Panel A. Within each subplot, the top row represents the distribution of Cronbach’s alphas (dark gray), and the following rows represent the distributions of correlations for RHA (red) and CHA (orange), respectively, plotted against the distribution of correlations for AA (light gray). Similar to Panel A, the top two subplots correspond to within-group predictions and the bottom two subplots correspond to across-group predictions.

**Figure S4.**
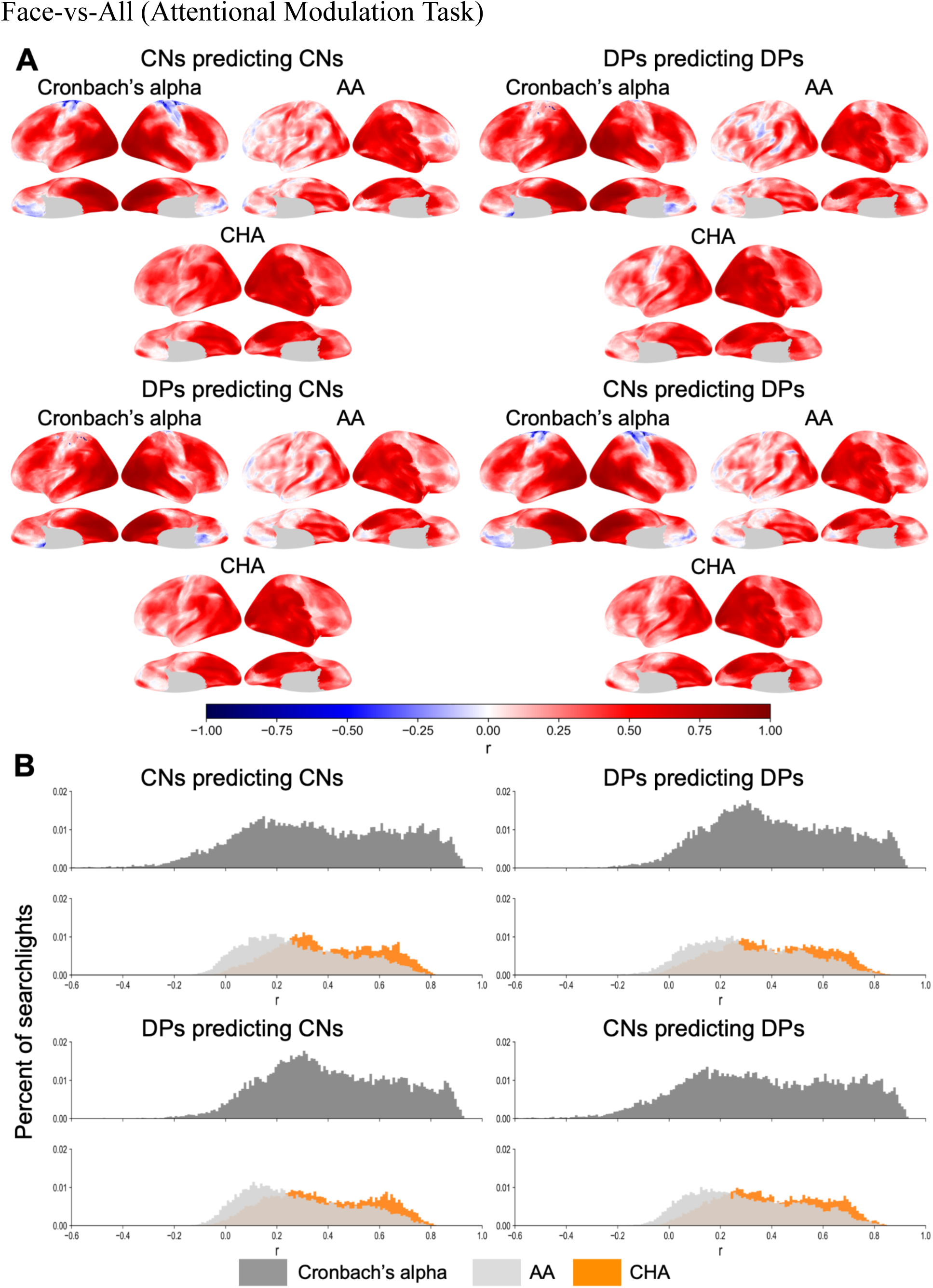
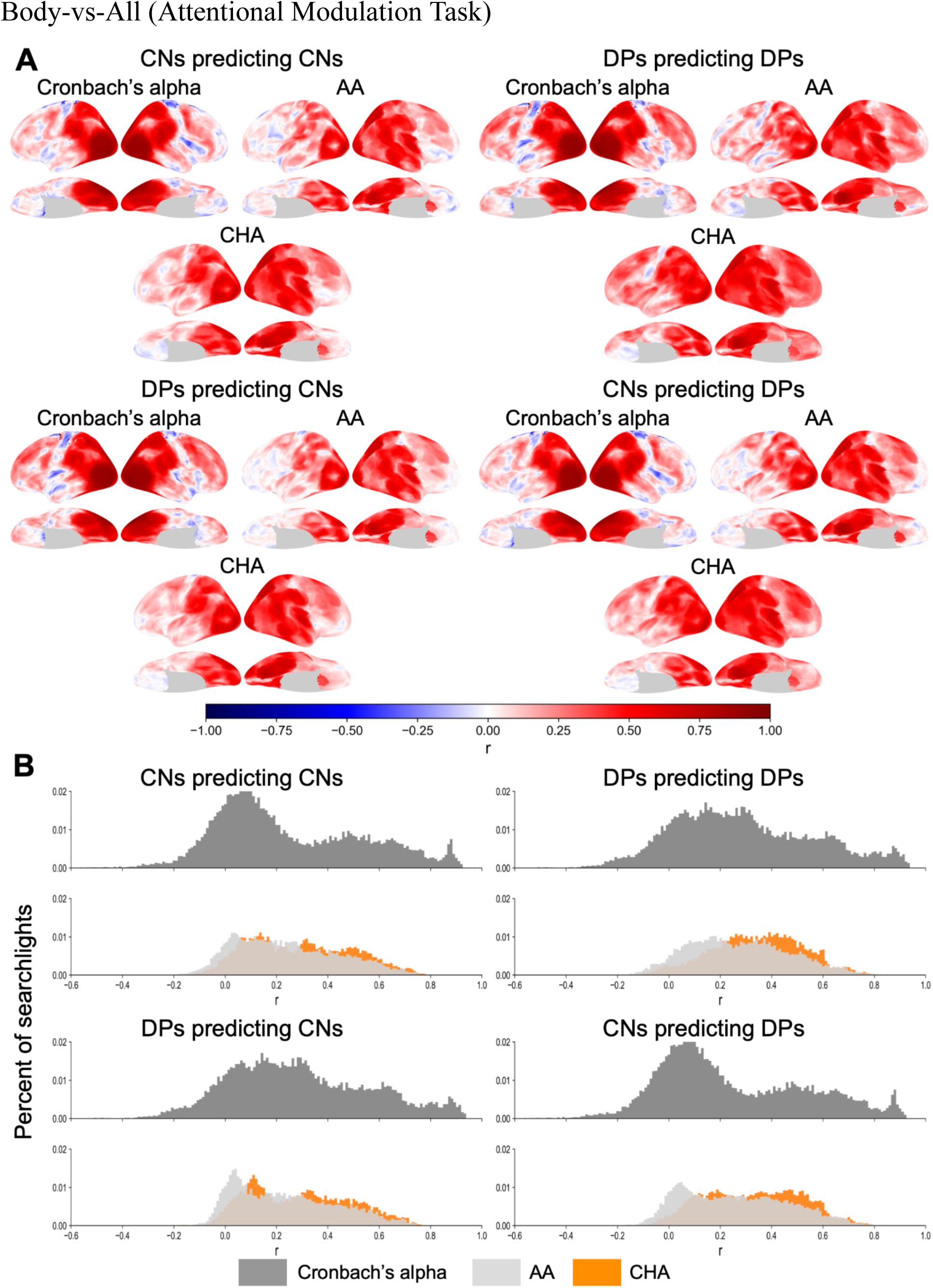
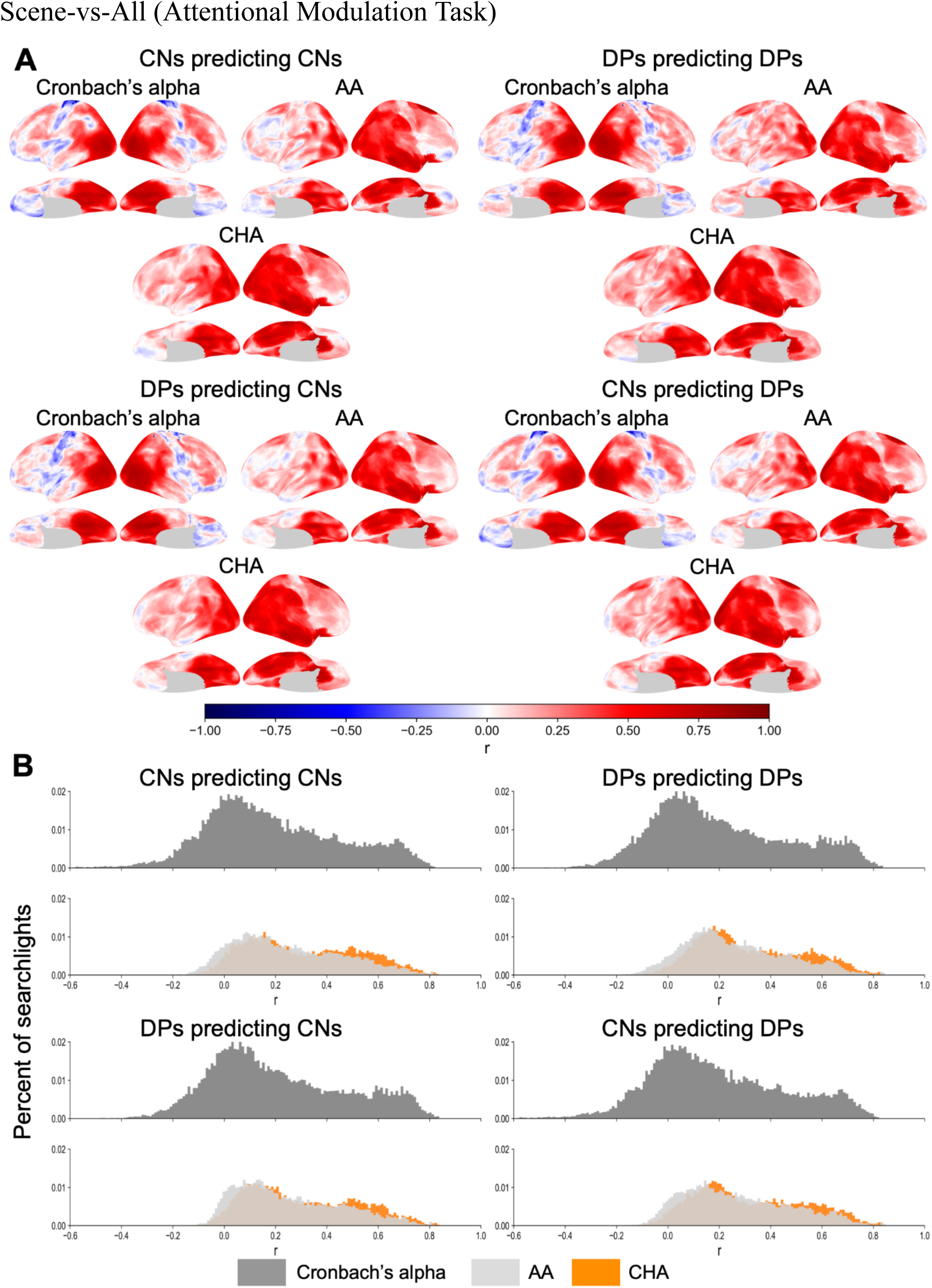
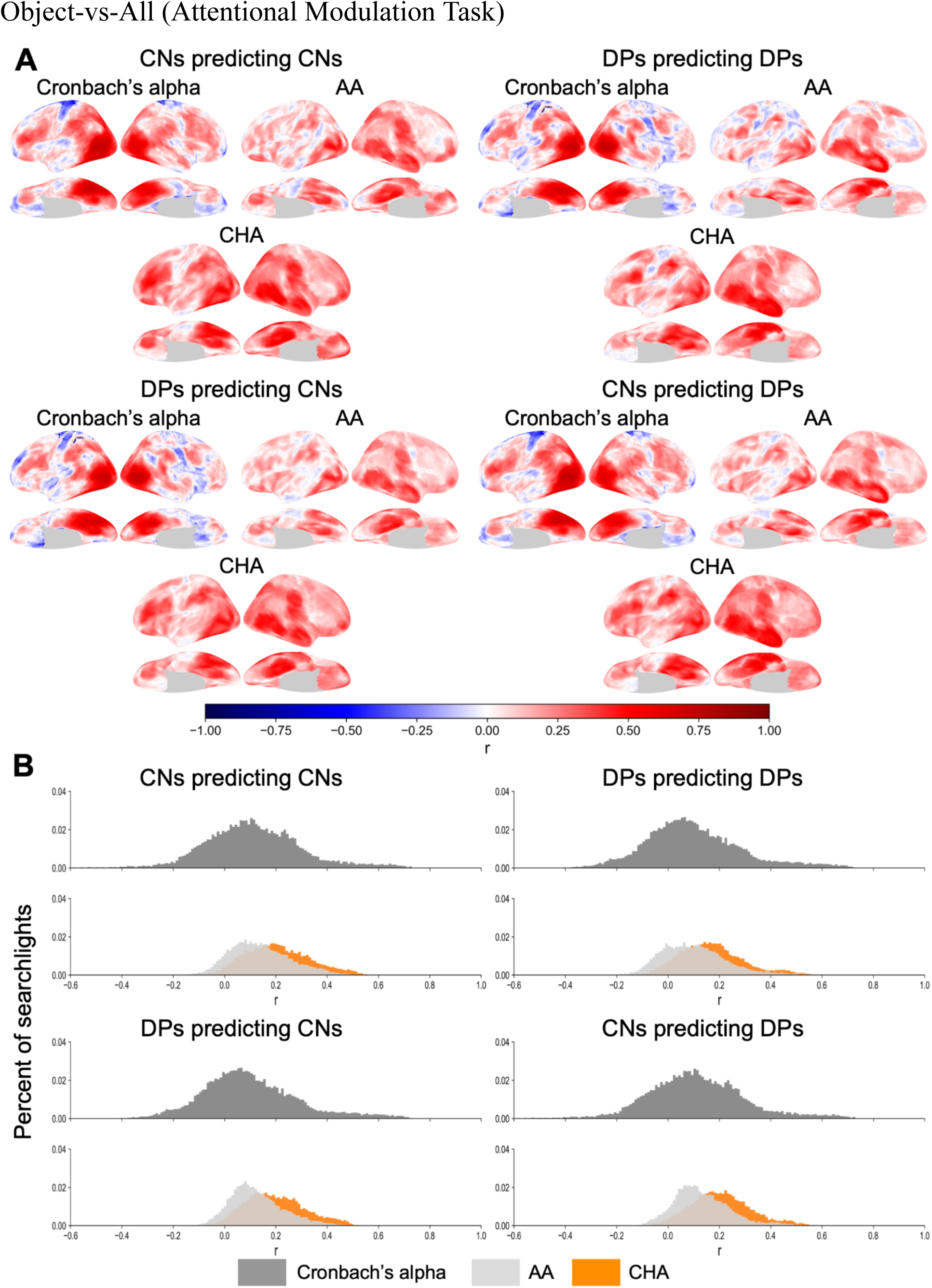
Searchlight analysis of Cronbach’s alpha and correlations for face-, body-, scene- and object-selective maps in the Dartmouth dataset using the attentional modulation task for hyperalignment. For each category, the contrast was defined as category vs. all other categories, and the plots follow the same layout as Figure 3 of the manuscript and Supplementary Figure S3.

**Figure S5.**
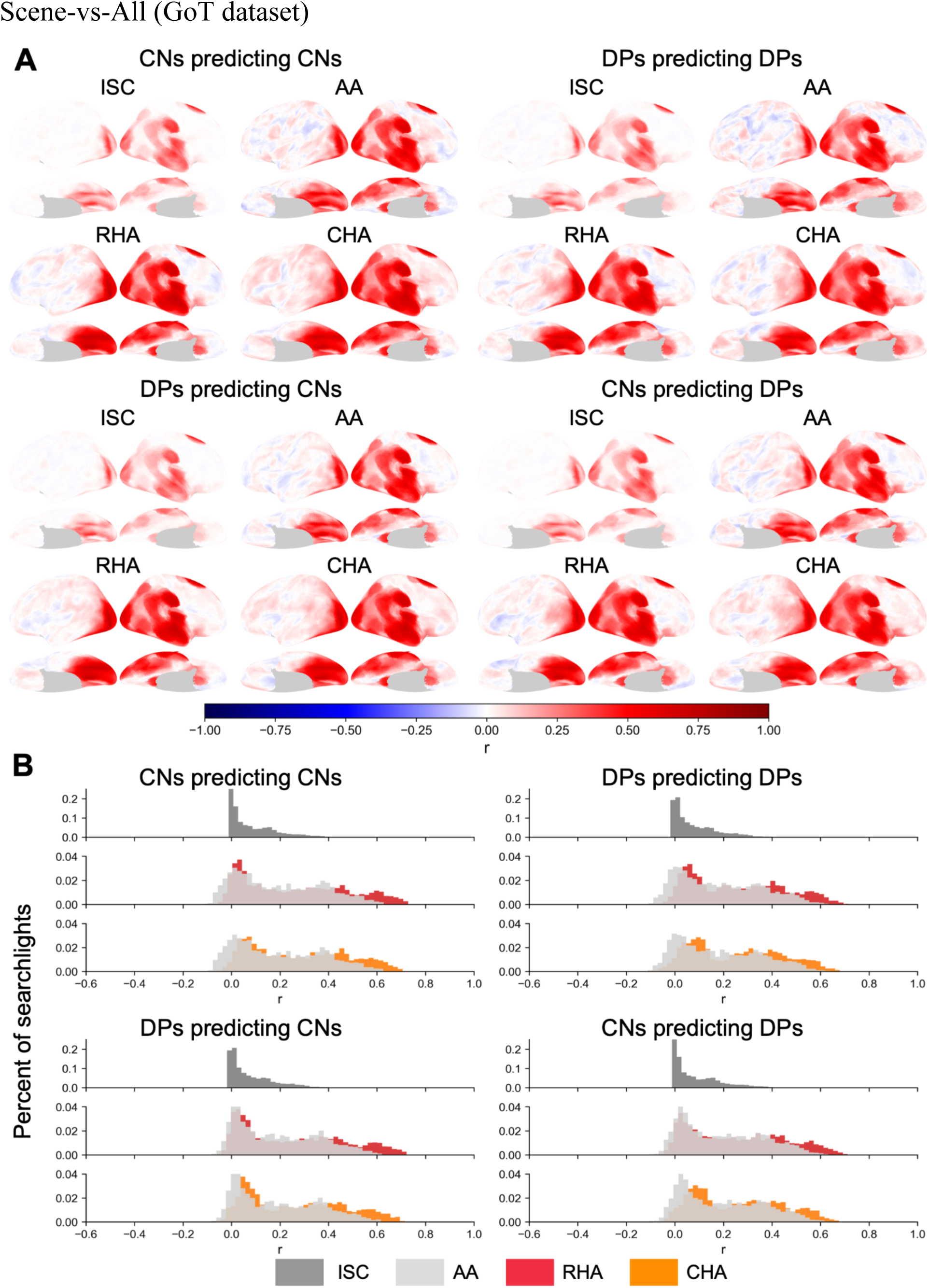
Searchlight analysis of Cronbach’s alpha and correlations for scene-selective maps in the GoT dataset. Scene-selective contrasts were calculated using scene vs all the other categories in the GoT localizers (faces, scrambled faces).

**Figure S6.**
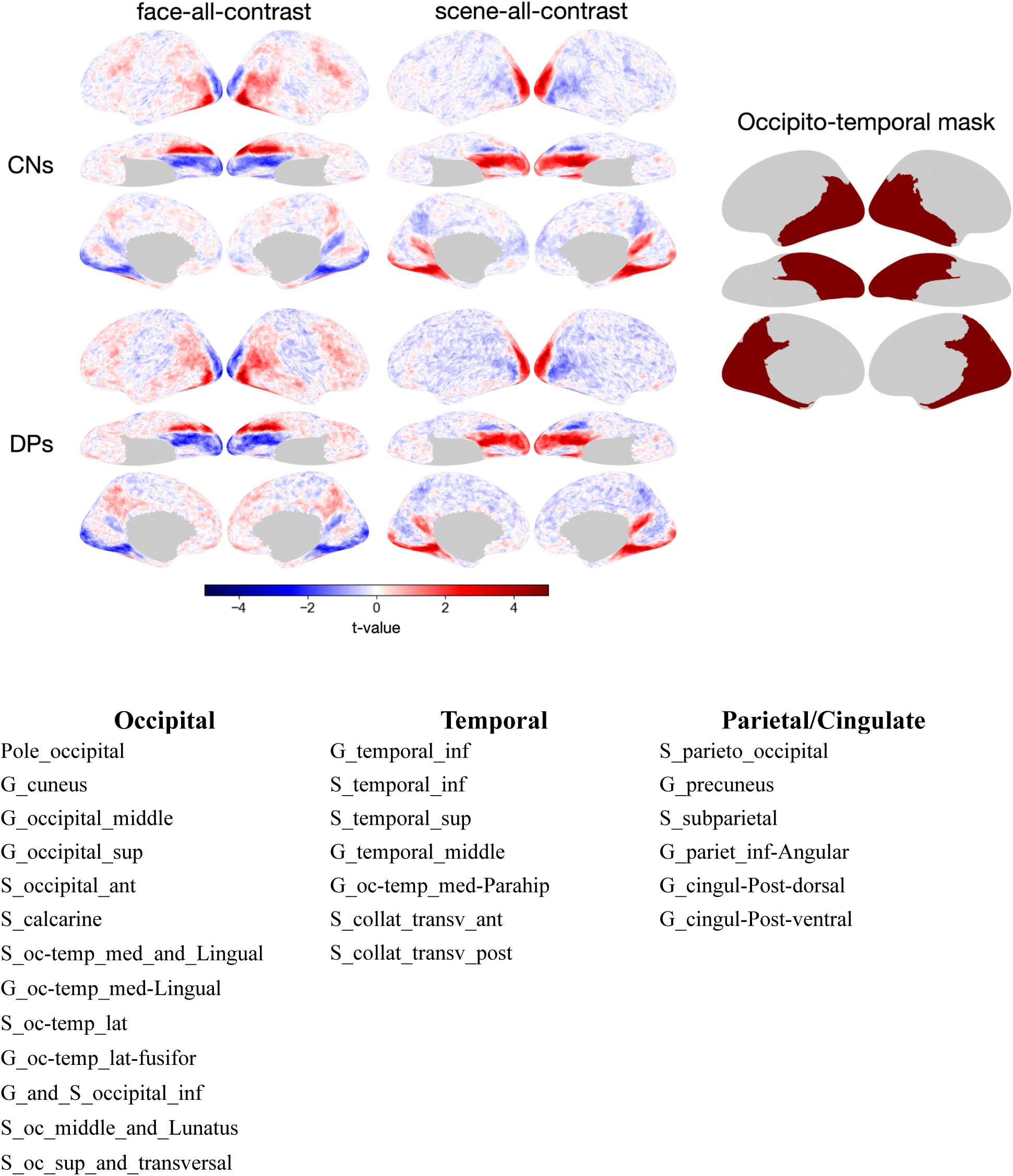
Occipito-temporal mask created using regions from the Desikan–Killiany parcellation (DK, aparc.a2009s) for the GoT dataset analysis. With a single run of static localizer, contrast maps estimated from participants’ own localizer data were noisier than the Dartmouth dataset, which had five dynamic localizer runs. With the GoT dataset, reliable activations were observed in occipito-temporal areas in both control and DP groups for both face and scene categories. To make a more robust comparison, we restricted our analysis of this dataset to the occipital-temporal areas using an anatomical mask derived from the DK parcellation. We selected regions from the DK parcellation to form a contiguous mask that broadly covers the occipital-temporal areas. The list shows the regions selected from the DK parcellation, in their original nomenclature, and corresponds to the masked areas in the right-most plot (the mask).

**Figure S7.**
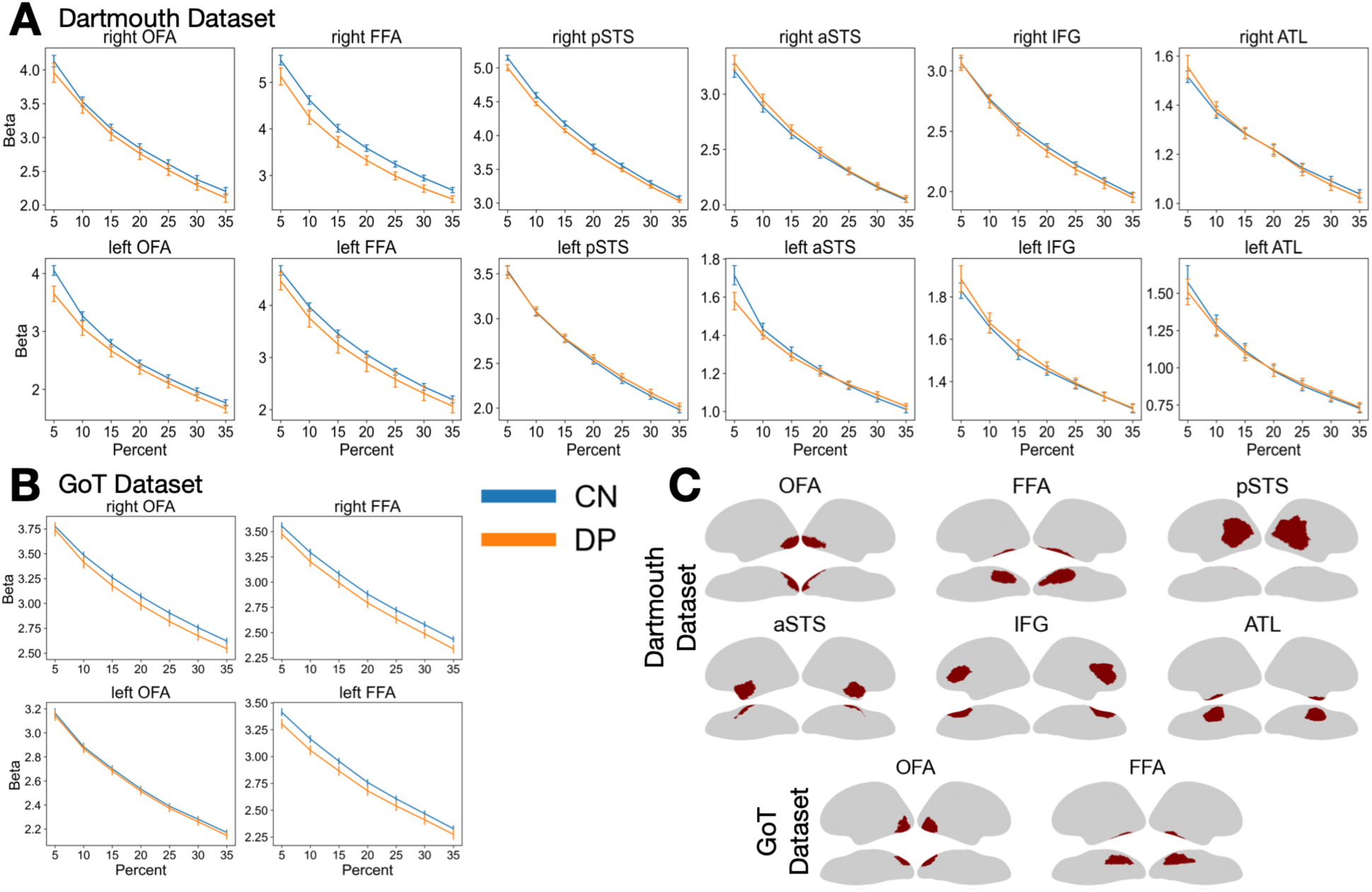
Comparison of face selectivity between controls and DPs in the face processing network (control: based on within-group predictions; DP: based on cross-group predictions). We define face selectivity here as the beta weights from the face vs. all other categories contrast for each participant, averaged in a region of interest (ROI). This figure complements the ROI analysis in the main paper, which compared face selectivity using CHA-predicted topographies from the within-group analysis (Controls-Controls, DP-DP). Here we used across-group predictions for the DP data (Controls predicting DPs) to conduct the ROI analysis. **(A)** Face selectivity differences in the Dartmouth dataset using localizer data predicted with CHA at ROI sizes from 5 to 35%. For each plot, the mean beta weight (y-axis) was calculated in a cross-validation procedure, by averaging four runs of the face vs. all contrast t-maps, selecting the top percent of voxels within a given ROI (x-axis), and averaging the beta weights within those top voxels in the left-out run in each of the five folds. **(B)** Group-level activation differences in the GoT dataset using localizer data predicted with CHA at ROI sizes from 5 to 35%. Two ventral ROIs were reliably localized in this dataset. Since there was only one run of localizer, the average beta weight was calculated by selecting the top X percent of voxels within that run’s t-map and extracting the corresponding beta weights. For both datasets, error bars indicate ±1 SE. **(C)** ROI masks for each dataset. The top two rows correspond to masks in the Dartmouth dataset, and the bottom row corresponds to masks in the GoT dataset.

**Figure S8.**
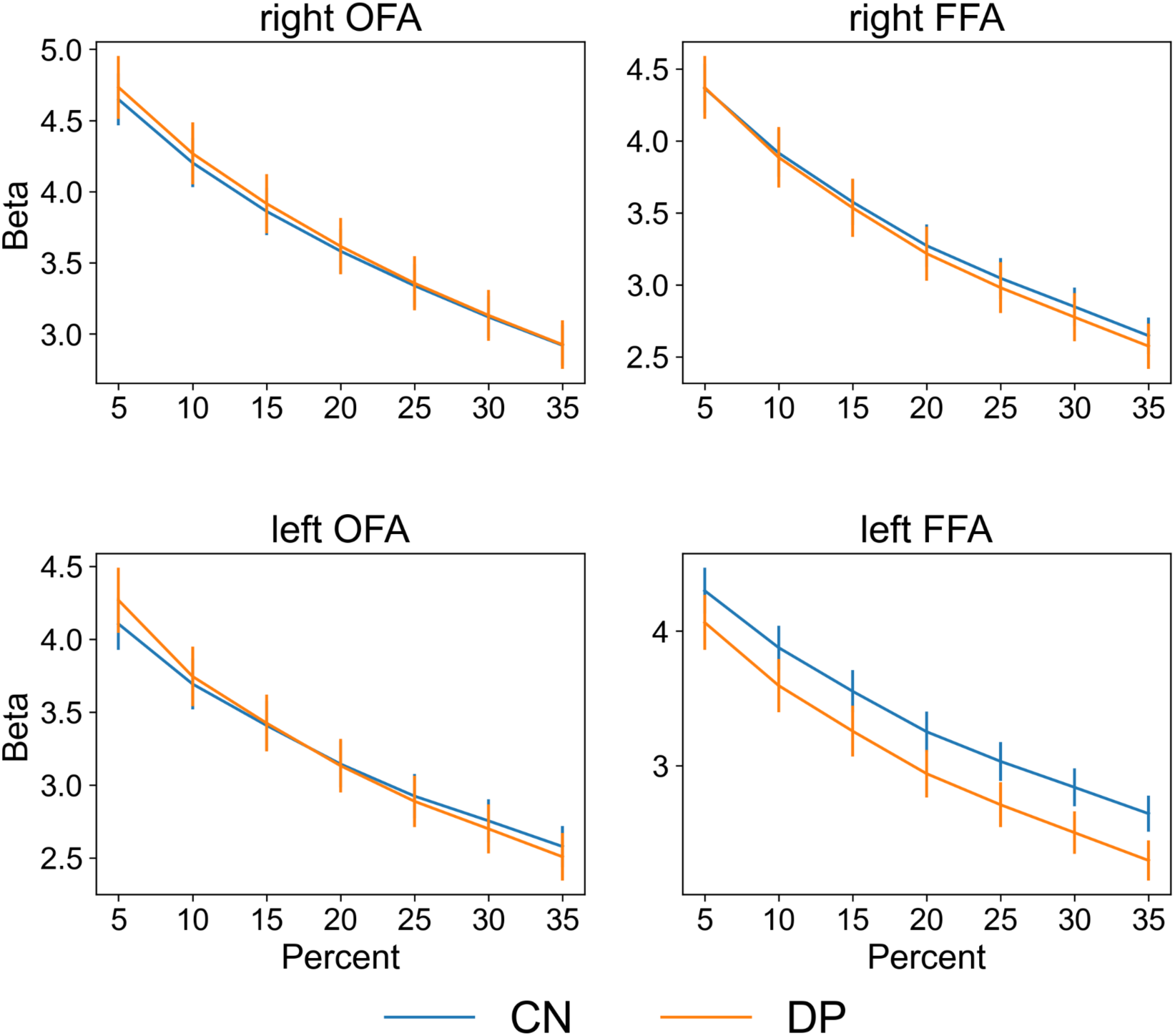
Comparison of face selectivity between controls and DPs in the face-selective ROIs in the GoT dataset using participants’ own localizer runs. Following the analysis steps in Jiahui et al., 2018, we manually prepared masks for the bilateral ROIs that generously cover possible vertices. Using 5% increments from 5 to 35%, we selected the top X% of voxels in these masks based on their t-values and extracted the beta weights for these selected voxels. Error bars indicate ±1 SE. No hyperalignment was involved in this analysis.

**Table 1.**
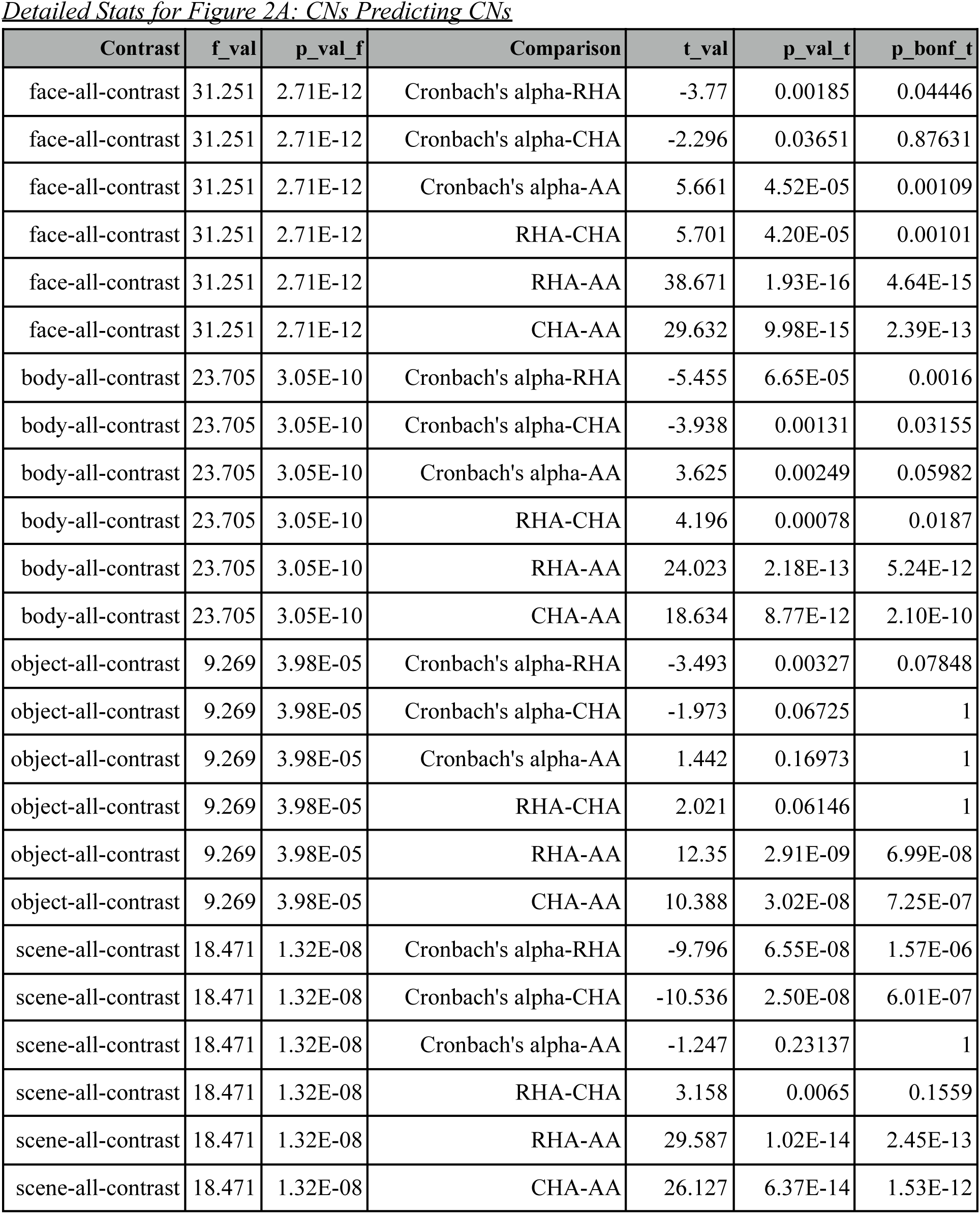
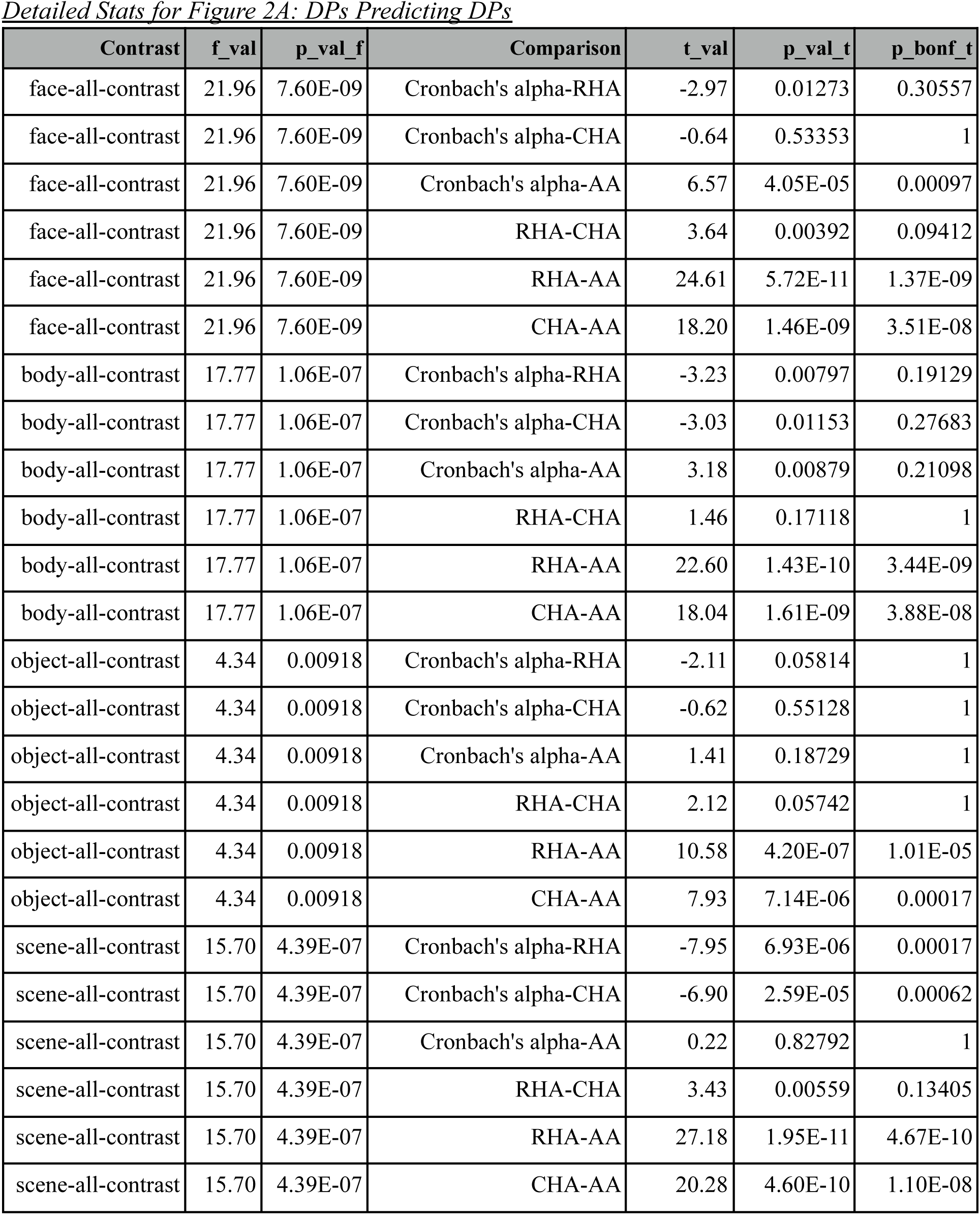
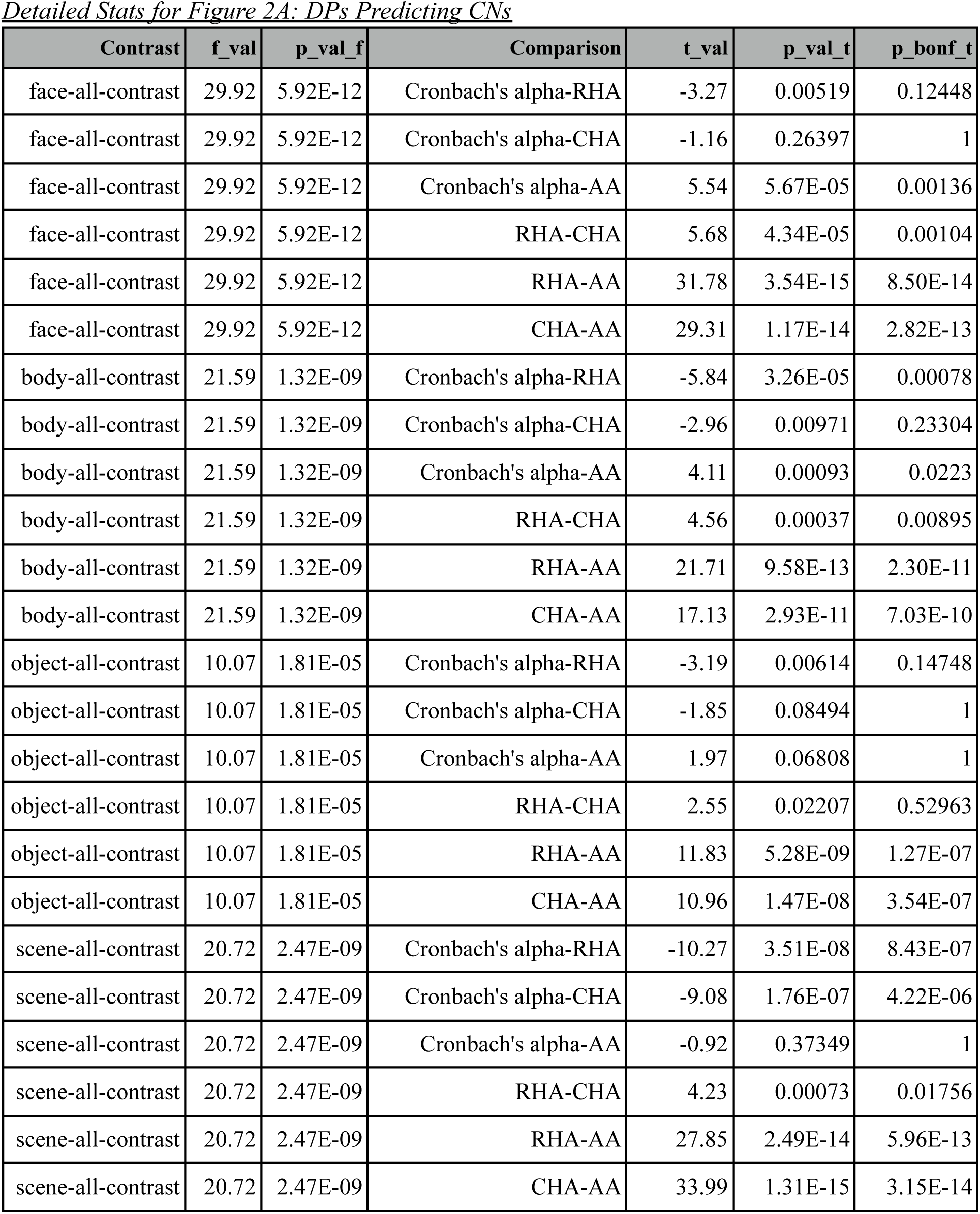
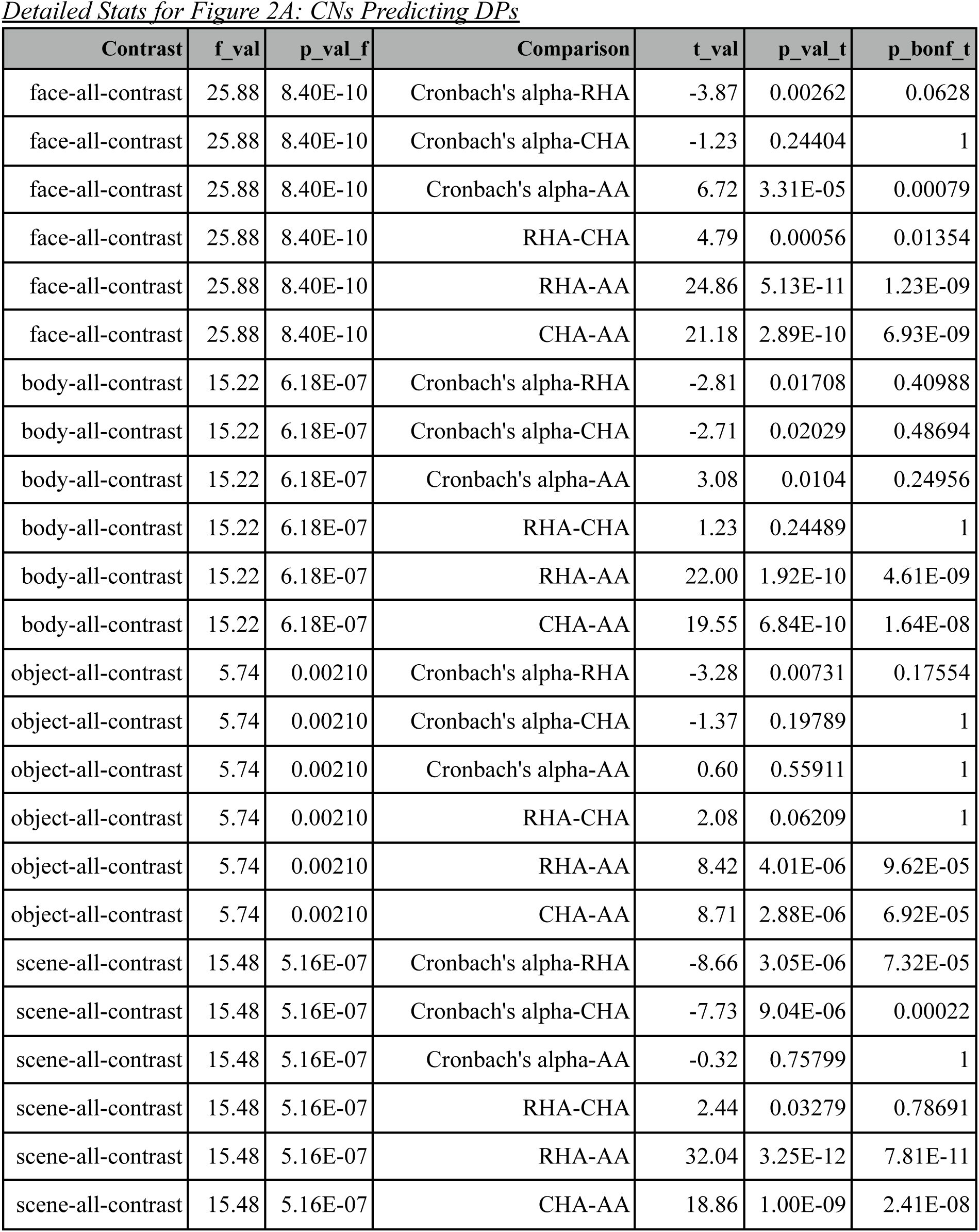
Statistical tables corresponding to the whole-brain analyses for the Dartmouth dataset (hyperalignment using the localizer task) in **Figure 2**. Each table corresponds to a barplot in the manuscript.

| Contrast | f_val | p_val_f | Comparison | t_val | p_val_t | p_bonf_t |
| --- | --- | --- | --- | --- | --- | --- |
| face-all-contrast | 31.251 | 2.71E-12 | Cronbach's alpha-RHA | -3.77 | 0.00185 | 0.04446 |
| face-all-contrast | 31.251 | 2.71E-12 | Cronbach's alpha-CHA | -2.296 | 0.03651 | 0.87631 |
| face-all-contrast | 31.251 | 2.71E-12 | Cronbach's alpha-AA | 5.661 | 4.52E-05 | 0.00109 |
| face-all-contrast | 31.251 | 2.71E-12 | RHA-CHA | 5.701 | 4.20E-05 | 0.00101 |
| face-all-contrast | 31.251 | 2.71E-12 | RHA-AA | 38.671 | 1.93E-16 | 4.64E-15 |
| face-all-contrast | 31.251 | 2.71E-12 | CHA-AA | 29.632 | 9.98E-15 | 2.39E-13 |
| body-all-contrast | 23.705 | 3.05E-10 | Cronbach's alpha-RHA | -5.455 | 6.65E-05 | 0.0016 |
| body-all-contrast | 23.705 | 3.05E-10 | Cronbach's alpha-CHA | -3.938 | 0.00131 | 0.03155 |
| body-all-contrast | 23.705 | 3.05E-10 | Cronbach's alpha-AA | 3.625 | 0.00249 | 0.05982 |
| body-all-contrast | 23.705 | 3.05E-10 | RHA-CHA | 4.196 | 0.00078 | 0.0187 |
| body-all-contrast | 23.705 | 3.05E-10 | RHA-AA | 24.023 | 2.18E-13 | 5.24E-12 |
| body-all-contrast | 23.705 | 3.05E-10 | CHA-AA | 18.634 | 8.77E-12 | 2.10E-10 |
| object-all-contrast | 9.269 | 3.98E-05 | Cronbach's alpha-RHA | -3.493 | 0.00327 | 0.07848 |
| object-all-contrast | 9.269 | 3.98E-05 | Cronbach's alpha-CHA | -1.973 | 0.06725 | 1 |
| object-all-contrast | 9.269 | 3.98E-05 | Cronbach's alpha-AA | 1.442 | 0.16973 | 1 |
| object-all-contrast | 9.269 | 3.98E-05 | RHA-CHA | 2.021 | 0.06146 | 1 |
| object-all-contrast | 9.269 | 3.98E-05 | RHA-AA | 12.35 | 2.91E-09 | 6.99E-08 |
| object-all-contrast | 9.269 | 3.98E-05 | CHA-AA | 10.388 | 3.02E-08 | 7.25E-07 |
| scene-all-contrast | 18.471 | 1.32E-08 | Cronbach's alpha-RHA | -9.796 | 6.55E-08 | 1.57E-06 |
| scene-all-contrast | 18.471 | 1.32E-08 | Cronbach's alpha-CHA | -10.536 | 2.50E-08 | 6.01E-07 |
| scene-all-contrast | 18.471 | 1.32E-08 | Cronbach's alpha-AA | -1.247 | 0.23137 | 1 |
| scene-all-contrast | 18.471 | 1.32E-08 | RHA-CHA | 3.158 | 0.0065 | 0.1559 |
| scene-all-contrast | 18.471 | 1.32E-08 | RHA-AA | 29.587 | 1.02E-14 | 2.45E-13 |
| scene-all-contrast | 18.471 | 1.32E-08 | CHA-AA | 26.127 | 6.37E-14 | 1.53E-12 |

*Detailed Stats for Figure 2A: DPs Predicting DPs*
| Contrast | f_val | p_val_f | Comparison | t_val | p_val_t | p_bonf_t |
| --- | --- | --- | --- | --- | --- | --- |
| face-all-contrast | 21.96 | 7.60E-09 | Cronbach's alpha-RHA | -2.97 | 0.01273 | 0.30557 |
| face-all-contrast | 21.96 | 7.60E-09 | Cronbach's alpha-CHA | -0.64 | 0.53353 | 1 |
| face-all-contrast | 21.96 | 7.60E-09 | Cronbach's alpha-AA | 6.57 | 4.05E-05 | 0.00097 |
| face-all-contrast | 21.96 | 7.60E-09 | RHA-CHA | 3.64 | 0.00392 | 0.09412 |
| face-all-contrast | 21.96 | 7.60E-09 | RHA-AA | 24.61 | 5.72E-11 | 1.37E-09 |
| face-all-contrast | 21.96 | 7.60E-09 | CHA-AA | 18.20 | 1.46E-09 | 3.51E-08 |
| body-all-contrast | 17.77 | 1.06E-07 | Cronbach's alpha-RHA | -3.23 | 0.00797 | 0.19129 |
| body-all-contrast | 17.77 | 1.06E-07 | Cronbach's alpha-CHA | -3.03 | 0.01153 | 0.27683 |
| body-all-contrast | 17.77 | 1.06E-07 | Cronbach's alpha-AA | 3.18 | 0.00879 | 0.21098 |
| body-all-contrast | 17.77 | 1.06E-07 | RHA-CHA | 1.46 | 0.17118 | 1 |
| body-all-contrast | 17.77 | 1.06E-07 | RHA-AA | 22.60 | 1.43E-10 | 3.44E-09 |
| body-all-contrast | 17.77 | 1.06E-07 | CHA-AA | 18.04 | 1.61E-09 | 3.88E-08 |
| object-all-contrast | 4.34 | 0.00918 | Cronbach's alpha-RHA | -2.11 | 0.05814 | 1 |
| object-all-contrast | 4.34 | 0.00918 | Cronbach's alpha-CHA | -0.62 | 0.55128 | 1 |
| object-all-contrast | 4.34 | 0.00918 | Cronbach's alpha-AA | 1.41 | 0.18729 | 1 |
| object-all-contrast | 4.34 | 0.00918 | RHA-CHA | 2.12 | 0.05742 | 1 |
| object-all-contrast | 4.34 | 0.00918 | RHA-AA | 10.58 | 4.20E-07 | 1.01E-05 |
| object-all-contrast | 4.34 | 0.00918 | CHA-AA | 7.93 | 7.14E-06 | 0.00017 |
| scene-all-contrast | 15.70 | 4.39E-07 | Cronbach's alpha-RHA | -7.95 | 6.93E-06 | 0.00017 |
| scene-all-contrast | 15.70 | 4.39E-07 | Cronbach's alpha-CHA | -6.90 | 2.59E-05 | 0.00062 |
| scene-all-contrast | 15.70 | 4.39E-07 | Cronbach's alpha-AA | 0.22 | 0.82792 | 1 |
| scene-all-contrast | 15.70 | 4.39E-07 | RHA-CHA | 3.43 | 0.00559 | 0.13405 |
| scene-all-contrast | 15.70 | 4.39E-07 | RHA-AA | 27.18 | 1.95E-11 | 4.67E-10 |
| scene-all-contrast | 15.70 | 4.39E-07 | CHA-AA | 20.28 | 4.60E-10 | 1.10E-08 |

*Detailed Stats for Figure 2A: DPs Predicting CNs*
| Contrast | f_val | p_val_f | Comparison | t_val | p_val_t | p_bonf_t |
| --- | --- | --- | --- | --- | --- | --- |
| face-all-contrast | 29.92 | 5.92E-12 | Cronbach's alpha-RHA | -3.27 | 0.00519 | 0.12448 |
| face-all-contrast | 29.92 | 5.92E-12 | Cronbach's alpha-CHA | -1.16 | 0.26397 | 1 |
| face-all-contrast | 29.92 | 5.92E-12 | Cronbach's alpha-AA | 5.54 | 5.67E-05 | 0.00136 |
| face-all-contrast | 29.92 | 5.92E-12 | RHA-CHA | 5.68 | 4.34E-05 | 0.00104 |
| face-all-contrast | 29.92 | 5.92E-12 | RHA-AA | 31.78 | 3.54E-15 | 8.50E-14 |
| face-all-contrast | 29.92 | 5.92E-12 | CHA-AA | 29.31 | 1.17E-14 | 2.82E-13 |
| body-all-contrast | 21.59 | 1.32E-09 | Cronbach's alpha-RHA | -5.84 | 3.26E-05 | 0.00078 |
| body-all-contrast | 21.59 | 1.32E-09 | Cronbach's alpha-CHA | -2.96 | 0.00971 | 0.23304 |
| body-all-contrast | 21.59 | 1.32E-09 | Cronbach's alpha-AA | 4.11 | 0.00093 | 0.0223 |
| body-all-contrast | 21.59 | 1.32E-09 | RHA-CHA | 4.56 | 0.00037 | 0.00895 |
| body-all-contrast | 21.59 | 1.32E-09 | RHA-AA | 21.71 | 9.58E-13 | 2.30E-11 |
| body-all-contrast | 21.59 | 1.32E-09 | CHA-AA | 17.13 | 2.93E-11 | 7.03E-10 |
| object-all-contrast | 10.07 | 1.81E-05 | Cronbach's alpha-RHA | -3.19 | 0.00614 | 0.14748 |
| object-all-contrast | 10.07 | 1.81E-05 | Cronbach's alpha-CHA | -1.85 | 0.08494 | 1 |
| object-all-contrast | 10.07 | 1.81E-05 | Cronbach's alpha-AA | 1.97 | 0.06808 | 1 |
| object-all-contrast | 10.07 | 1.81E-05 | RHA-CHA | 2.55 | 0.02207 | 0.52963 |
| object-all-contrast | 10.07 | 1.81E-05 | RHA-AA | 11.83 | 5.28E-09 | 1.27E-07 |
| object-all-contrast | 10.07 | 1.81E-05 | CHA-AA | 10.96 | 1.47E-08 | 3.54E-07 |
| scene-all-contrast | 20.72 | 2.47E-09 | Cronbach's alpha-RHA | -10.27 | 3.51E-08 | 8.43E-07 |
| scene-all-contrast | 20.72 | 2.47E-09 | Cronbach's alpha-CHA | -9.08 | 1.76E-07 | 4.22E-06 |
| scene-all-contrast | 20.72 | 2.47E-09 | Cronbach's alpha-AA | -0.92 | 0.37349 | 1 |
| scene-all-contrast | 20.72 | 2.47E-09 | RHA-CHA | 4.23 | 0.00073 | 0.01756 |
| scene-all-contrast | 20.72 | 2.47E-09 | RHA-AA | 27.85 | 2.49E-14 | 5.96E-13 |
| scene-all-contrast | 20.72 | 2.47E-09 | CHA-AA | 33.99 | 1.31E-15 | 3.15E-14 |

*Detailed Stats for Figure 2A: CNs Predicting DPs*
| Contrast | f_val | p_val_f | Comparison | t_val | p_val_t | p_bonf_t |
| --- | --- | --- | --- | --- | --- | --- |
| face-all-contrast | 25.88 | 8.40E-10 | Cronbach's alpha-RHA | -3.87 | 0.00262 | 0.0628 |
| face-all-contrast | 25.88 | 8.40E-10 | Cronbach's alpha-CHA | -1.23 | 0.24404 | 1 |
| face-all-contrast | 25.88 | 8.40E-10 | Cronbach's alpha-AA | 6.72 | 3.31E-05 | 0.00079 |
| face-all-contrast | 25.88 | 8.40E-10 | RHA-CHA | 4.79 | 0.00056 | 0.01354 |
| face-all-contrast | 25.88 | 8.40E-10 | RHA-AA | 24.86 | 5.13E-11 | 1.23E-09 |
| face-all-contrast | 25.88 | 8.40E-10 | CHA-AA | 21.18 | 2.89E-10 | 6.93E-09 |
| body-all-contrast | 15.22 | 6.18E-07 | Cronbach's alpha-RHA | -2.81 | 0.01708 | 0.40988 |
| body-all-contrast | 15.22 | 6.18E-07 | Cronbach's alpha-CHA | -2.71 | 0.02029 | 0.48694 |
| body-all-contrast | 15.22 | 6.18E-07 | Cronbach's alpha-AA | 3.08 | 0.0104 | 0.24956 |
| body-all-contrast | 15.22 | 6.18E-07 | RHA-CHA | 1.23 | 0.24489 | 1 |
| body-all-contrast | 15.22 | 6.18E-07 | RHA-AA | 22.00 | 1.92E-10 | 4.61E-09 |
| body-all-contrast | 15.22 | 6.18E-07 | CHA-AA | 19.55 | 6.84E-10 | 1.64E-08 |
| object-all-contrast | 5.74 | 0.00210 | Cronbach's alpha-RHA | -3.28 | 0.00731 | 0.17554 |
| object-all-contrast | 5.74 | 0.00210 | Cronbach's alpha-CHA | -1.37 | 0.19789 | 1 |
| object-all-contrast | 5.74 | 0.00210 | Cronbach's alpha-AA | 0.60 | 0.55911 | 1 |
| object-all-contrast | 5.74 | 0.00210 | RHA-CHA | 2.08 | 0.06209 | 1 |
| object-all-contrast | 5.74 | 0.00210 | RHA-AA | 8.42 | 4.01E-06 | 9.62E-05 |
| object-all-contrast | 5.74 | 0.00210 | CHA-AA | 8.71 | 2.88E-06 | 6.92E-05 |
| scene-all-contrast | 15.48 | 5.16E-07 | Cronbach's alpha-RHA | -8.66 | 3.05E-06 | 7.32E-05 |
| scene-all-contrast | 15.48 | 5.16E-07 | Cronbach's alpha-CHA | -7.73 | 9.04E-06 | 0.00022 |
| scene-all-contrast | 15.48 | 5.16E-07 | Cronbach's alpha-AA | -0.32 | 0.75799 | 1 |
| scene-all-contrast | 15.48 | 5.16E-07 | RHA-CHA | 2.44 | 0.03279 | 0.78691 |
| scene-all-contrast | 15.48 | 5.16E-07 | RHA-AA | 32.04 | 3.25E-12 | 7.81E-11 |
| scene-all-contrast | 15.48 | 5.16E-07 | CHA-AA | 18.86 | 1.00E-09 | 2.41E-08 |

**Table 2.** Statistical tables corresponding to the whole-brain analyses for the Dartmouth dataset (hyperalignment using the attentional modulation task) in **Figure S2**. Each table corresponds to a barplot, and the previously-described methods were followed here.

| Contrast | f_val | p_val_f | Comparison | t_val | p_val_t | p_bonf_t |
| --- | --- | --- | --- | --- | --- | --- |
| face-all-contrast | 12.087 | 6.29E-05 | Cronbach's alpha-CHA | 3.234 | 5.57E-03 | 6.68E-02 |
| face-all-contrast | 12.087 | 6.29E-05 | Cronbach's alpha-AA | 5.661 | 4.52E-05 | 0.00054 |
| face-all-contrast | 12.087 | 6.29E-05 | CHA-AA | 7.462 | 2.01E-06 | 2.41E-05 |
| body-all-contrast | 3.893 | 2.76E-02 | Cronbach's alpha-CHA | 2.007 | 6.31E-02 | 7.58E-01 |
| body-all-contrast | 3.893 | 2.76E-02 | Cronbach's alpha-AA | 3.625 | 2.49E-03 | 2.99E-02 |
| body-all-contrast | 3.893 | 2.76E-02 | CHA-AA | 5.063 | 1.40E-04 | 1.68E-03 |
| object-all-contrast | 1.81 | 1.75E-01 | Cronbach's alpha-CHA | -0.25 | 8.06E-01 | 1 |
| object-all-contrast | 1.81 | 1.75E-01 | Cronbach's alpha-AA | 1.442 | 0.16973 | 1 |
| object-all-contrast | 1.81 | 1.75E-01 | CHA-AA | 6.426 | 1.14E-05 | 0.00014 |
| scene-all-contrast | 1.975 | 1.51E-01 | Cronbach's alpha-CHA | -3.969 | 1.24E-03 | 1.48E-02 |
| scene-all-contrast | 1.975 | 1.51E-01 | Cronbach's alpha-AA | -1.247 | 2.31E-01 | 1 |
| scene-all-contrast | 1.975 | 1.51E-01 | CHA-AA | 8.515 | 3.96E-07 | 4.75E-06 |

*Detailed Stats for Figure S2A: DPs Predicting DPs*
| Contrast | f_val | p_val_f | Comparison | t_val | p_val_t | p_bonf_t |
| --- | --- | --- | --- | --- | --- | --- |
| face-all-contrast | 15.5 | 1.79E-05 | Cronbach's alpha-CHA | 3.762 | 0.00314 | 0.03773 |
| face-all-contrast | 15.5 | 1.79E-05 | Cronbach's alpha-AA | 6.565 | 4.05E-05 | 0.00049 |
| face-all-contrast | 15.5 | 1.79E-05 | CHA-AA | 12.293 | 9.07E-08 | 1.09E-06 |
| body-all-contrast | 4.639 | 0.01678 | Cronbach's alpha-CHA | 2.021 | 0.06835 | 0.82021 |
| body-all-contrast | 4.639 | 0.01678 | Cronbach's alpha-AA | 3.178 | 0.00879 | 0.10549 |
| body-all-contrast | 4.639 | 0.01678 | CHA-AA | 3.725 | 0.00335 | 0.04022 |
| object-all-contrast | 1.287 | 0.28971 | Cronbach's alpha-CHA | 0.135 | 0.89523 | 1 |
| object-all-contrast | 1.287 | 0.28971 | Cronbach's alpha-AA | 1.406 | 0.18729 | 1 |
| object-all-contrast | 1.287 | 0.28971 | CHA-AA | 5.826 | 0.00011 | 0.00138 |
| scene-all-contrast | 1.433 | 0.25309 | Cronbach's alpha-CHA | -2.035 | 0.06668 | 0.80017 |
| scene-all-contrast | 1.433 | 0.25309 | Cronbach's alpha-AA | 0.223 | 0.82792 | 1 |
| scene-all-contrast | 1.433 | 0.25309 | CHA-AA | 11.648 | 1.58E-07 | 1.89E-06 |

*Detailed Stats for Figure S2A: DPs Predicting CNs*
| Contrast | f_val | p_val_f | Comparison | t_val | p_val_t | p_bonf_t |
| --- | --- | --- | --- | --- | --- | --- |
| face-all-contrast | 15.352 | 1.94E-05 | Cronbach's alpha-CHA | 4.345 | 0.00117 | 0.01398 |
| face-all-contrast | 15.352 | 1.94E-05 | Cronbach's alpha-AA | 6.715 | 3.31E-05 | 0.0004 |
| face-all-contrast | 15.352 | 1.94E-05 | CHA-AA | 6.787 | 3.01E-05 | 0.00036 |
| body-all-contrast | 4.496 | 0.01875 | Cronbach's alpha-CHA | 2.225 | 0.04797 | 0.57561 |
| body-all-contrast | 4.496 | 0.01875 | Cronbach's alpha-AA | 3.084 | 0.0104 | 0.12478 |
| body-all-contrast | 4.496 | 0.01875 | CHA-AA | 3.622 | 0.00401 | 0.04814 |
| object-all-contrast | 0.894 | 0.41889 | Cronbach's alpha-CHA | -0.647 | 0.53071 | 1 |
| object-all-contrast | 0.894 | 0.41889 | Cronbach's alpha-AA | 0.602 | 0.55911 | 1 |
| object-all-contrast | 0.894 | 0.41889 | CHA-AA | 5.078 | 0.00036 | 0.00427 |
| scene-all-contrast | 1.24 | 0.30259 | Cronbach's alpha-CHA | -2.258 | 0.04528 | 0.54333 |
| scene-all-contrast | 1.24 | 0.30259 | Cronbach's alpha-AA | -0.316 | 0.75799 | 1 |
| scene-all-contrast | 1.24 | 0.30259 | CHA-AA | 10.732 | 3.63E-07 | 4.36E-06 |

*Detailed Stats for Figure S2A: CNs Predicting DPs*
| Contrast | f_val | p_val_f | Comparison | t_val | p_val_t | p_bonf_t |
| --- | --- | --- | --- | --- | --- | --- |
| face-all-contrast | 13.783 | 2.14E-05 | Cronbach's alpha-CHA | 3.14 | 0.00675 | 0.08097 |
| face-all-contrast | 13.783 | 2.14E-05 | Cronbach's alpha-AA | 5.539 | 5.67E-05 | 0.00068 |
| face-all-contrast | 13.783 | 2.14E-05 | CHA-AA | 9.766 | 6.82E-08 | 8.18E-07 |
| body-all-contrast | 4.474 | 0.0169 | Cronbach's alpha-CHA | 2.987 | 0.00922 | 0.11059 |
| body-all-contrast | 4.474 | 0.0169 | Cronbach's alpha-AA | 4.109 | 0.00093 | 0.01115 |
| body-all-contrast | 4.474 | 0.0169 | CHA-AA | 3.68 | 0.00223 | 0.02672 |
| object-all-contrast | 2.873 | 0.06698 | Cronbach's alpha-CHA | -0.144 | 0.8872 | 1 |
| object-all-contrast | 2.873 | 0.06698 | Cronbach's alpha-AA | 1.966 | 0.06808 | 0.81697 |
| object-all-contrast | 2.873 | 0.06698 | CHA-AA | 6.856 | 5.45E-06 | 6.55E-05 |
| scene-all-contrast | 1.806 | 0.17598 | Cronbach's alpha-CHA | -3.213 | 0.00581 | 0.06968 |
| scene-all-contrast | 1.806 | 0.17598 | Cronbach's alpha-AA | -0.917 | 0.37349 | 1 |
| scene-all-contrast | 1.806 | 0.17598 | CHA-AA | 9.836 | 6.21E-08 | 7.45E-07 |

**Table 3.**
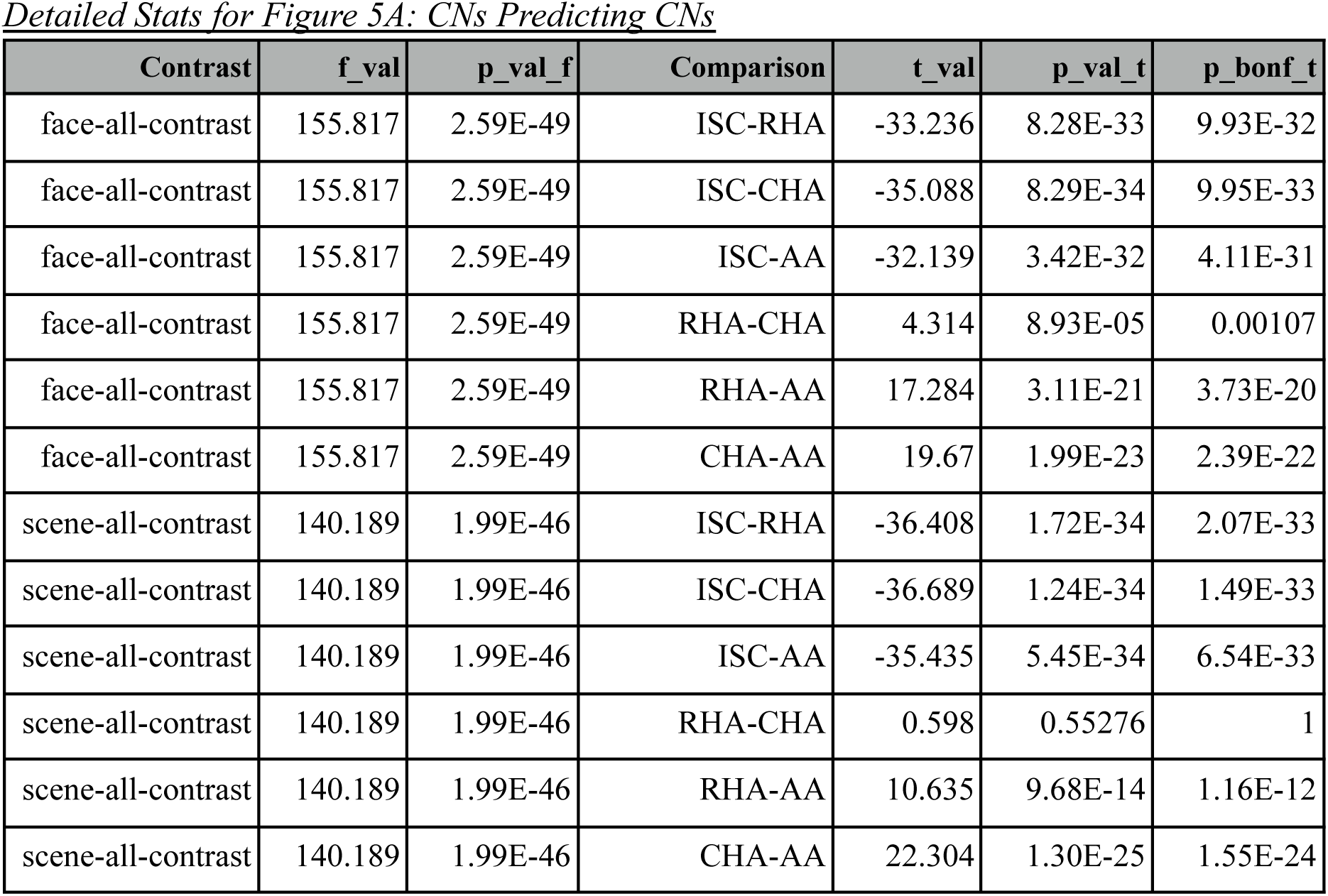
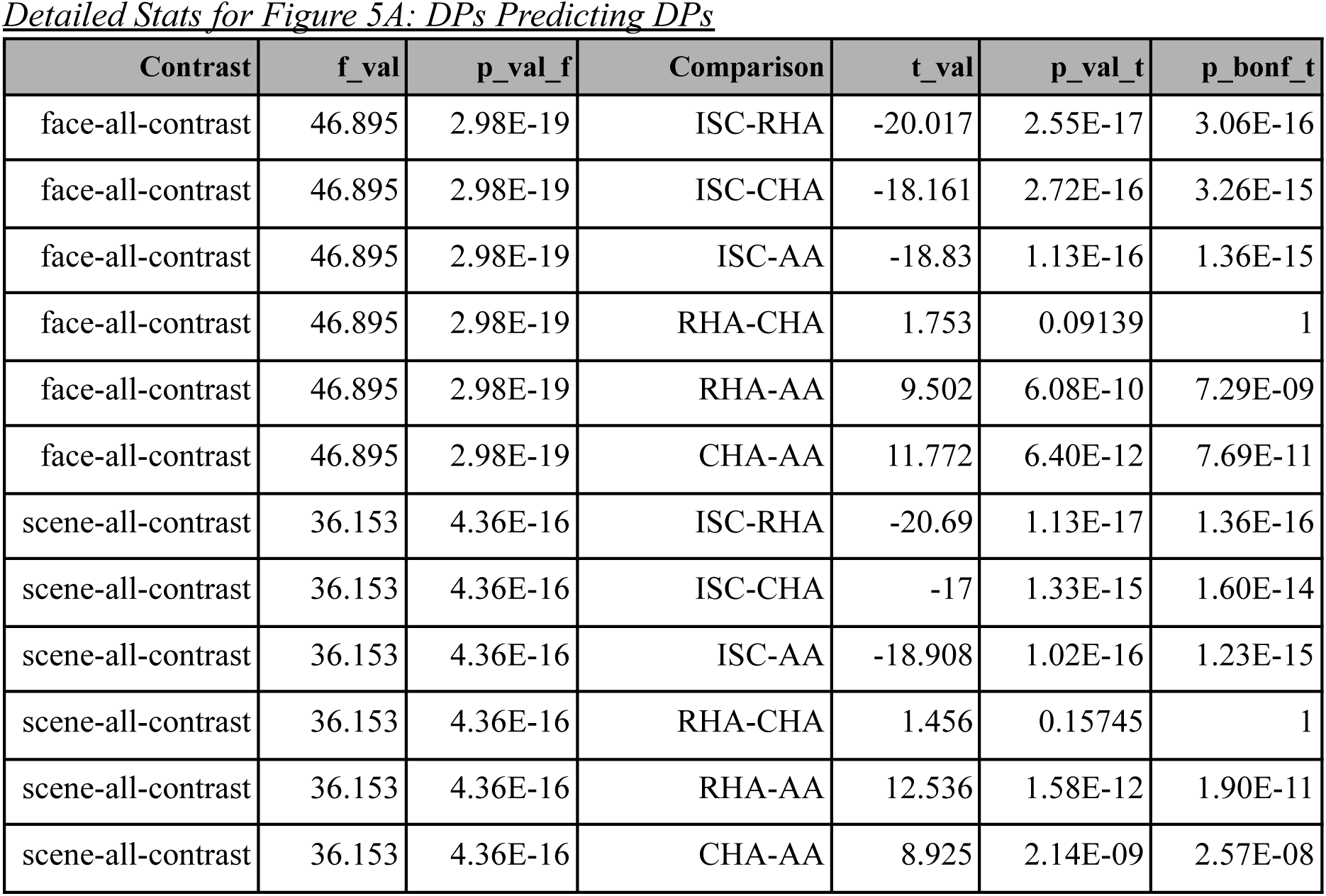
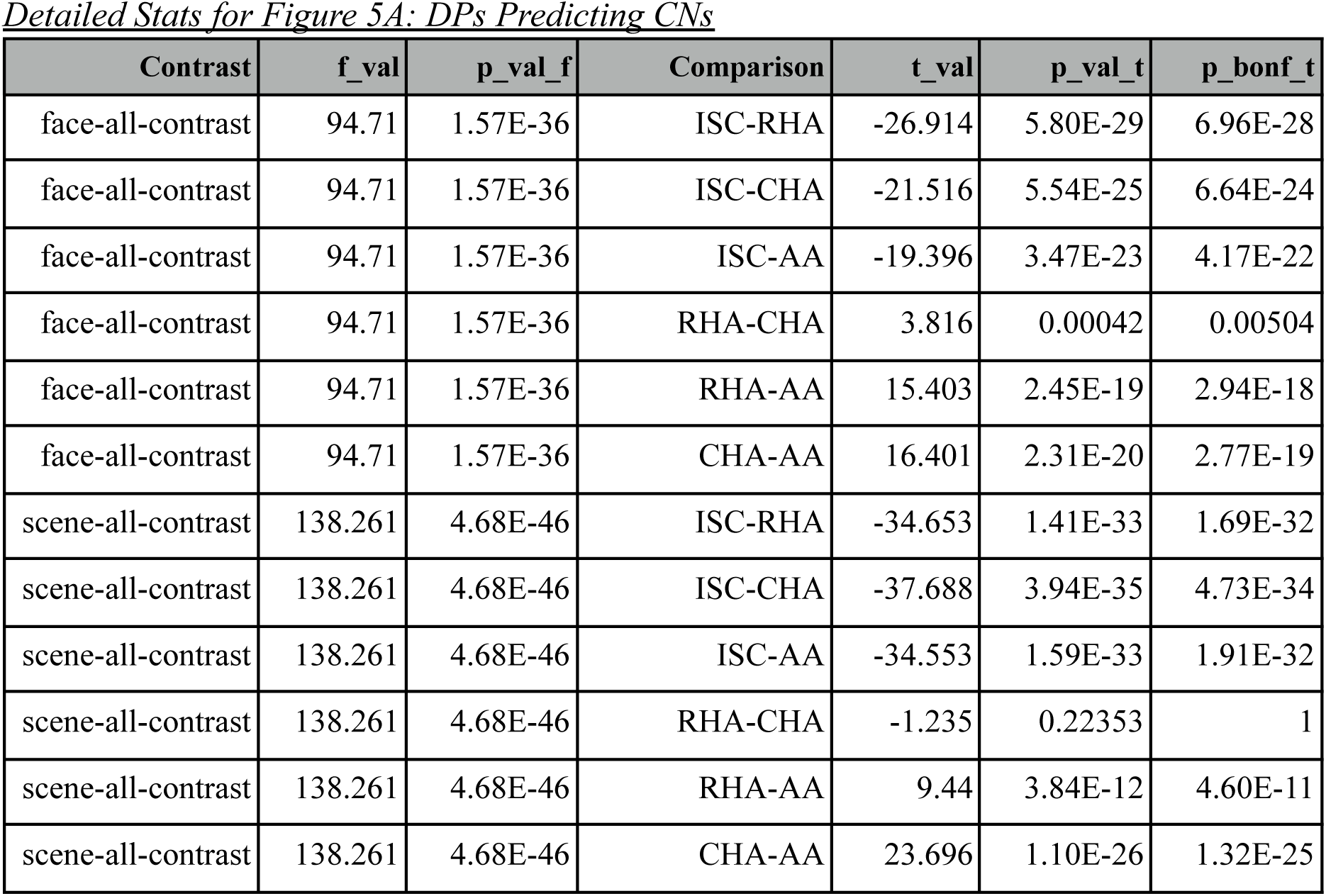
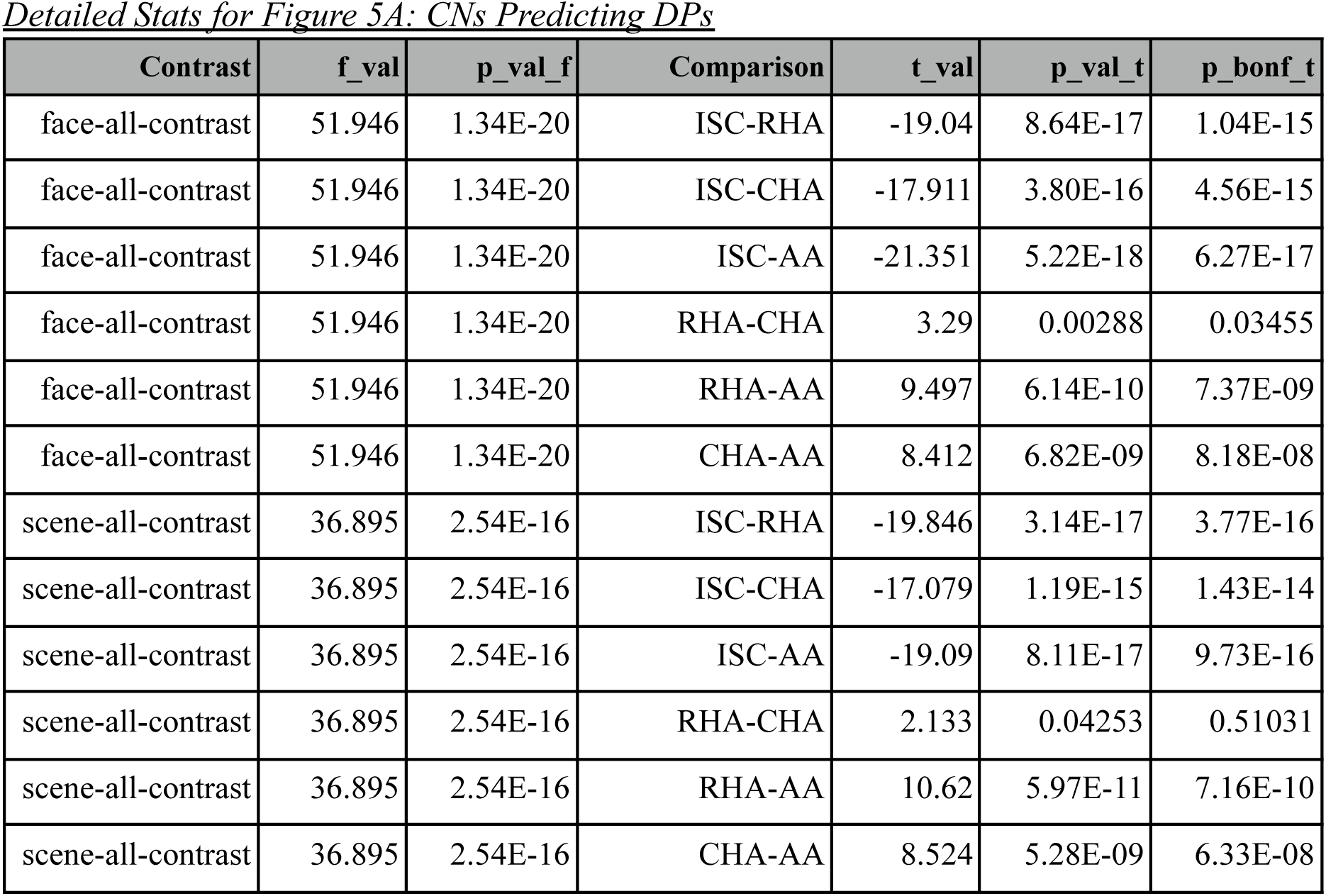
Statistical tables corresponding to the whole-brain analyses for the GoT dataset in **Figure 5**. Each table corresponds to a barplot in the manuscript.

